# Chronic warming, not heatwaves, predicts tropical fish colonization in the Mediterranean

**DOI:** 10.64898/2026.09.03.749106

**Authors:** Shahar Chaikin, Georgios Vagenas, Miguel Matias, Marika Galanidi, Argyro Zenetos, Miguel B. Araújo

## Abstract

The opening of the Suez Canal abruptly reconnected the Red Sea and the Mediterranean Sea, collapsing a geological barrier that had separated their biotas for millions of years. This exchange offers a natural experiment to determine whether tropical species’ spread at their cold-edge range is driven by acute thermal shocks (marine heatwaves) or by progressive easing of physiological constraints under sustained warming. We analyzed 1,059 first-arrival time series for 75 non-indigenous fish species across the Mediterranean, explicitly accounting for progressive salinification, oceanographic connectivity, and historical canal enlargement. Our models demonstrate that colonization dynamics are overwhelmingly dominated by ecological momentum—the time-dependent geographic spread of populations—and facilitated by chronic background warming. Marine heatwaves explain negligible variance and show no detectable effect on colonization success. Instead, warming operates as a stage-dependent continuous climatic facilitator, accelerating post-establishment expansion, rather than triggering introductions. Thus, this shift toward warm-adapted communities is governed by chronic warming, with the Suez Canal triggering the reorganization of Mediterranean biogeography.

## Introduction

The construction of the Suez Canal initiated one of the most consequential biogeographical experiments of the Anthropocene, effectively reversing the geological separation of the Red Sea and Mediterranean biotas [1]. By creating a permanent maritime corridor, it enabled a strongly asymmetric flow of hundreds of tropical and subtropical Indo-Pacific species—often called Lessepsian migrants—into the temperate Mediterranean Sea. This anthropogenic reconnection is a maritime analogue of the natural formation of the Isthmus of Panama, which joined the Americas and triggered a major terrestrial biotic interchange [2]. As Lessepsian fishes continue to expand and establish across the Mediterranean [3], they provide a uniquely resolved opportunity to disentangle the mechanisms of climate-driven redistribution. Specifically, this “natural laboratory” allows us to address a central uncertainty in climate-change ecology: are the poleward range shifts in marine species [4–10] driven primarily by episodic thermal extremes, or by the gradual relaxation of physiological constraints under chronic warming?

Extreme weather events, such as Marine Heatwaves (MHWs), are hypothesized to facilitate colonization by opening short-lived “windows of opportunity” [11–15], for example, by temporarily weakening biotic resistance in native communities [16] or creating instantaneous thermal or spatial bridges for warm-affinity species [17,18]. However, empirical support for this “MHW-driven” mechanism remains equivocal. While some studies report that MHWs favor non-indigenous range-expanding species [19,20], others find that thermal extremes negatively impact these populations [21–23], or contribute little relative to long-term trends [24]. This conflict underscores a persistent gap: we still lack consensus on whether biodiversity redistribution is governed by episodic and extreme events or by the chronic upward drift of temperature baselines.

Quantifying the specific role of climate in this redistribution of species requires separating climatic forcing from ecological momentum, i.e., the intrinsic, time-dependent expansion of populations driven by introduction history and ongoing spread [25]. In the Mediterranean, such momentum may reflect the cumulative effects of propagule pressure, species traits, and positive feedbacks associated with an open invasion corridor, such as the Suez Canal [26–29]. Together, these processes can generate a background rate of spread that is partly independent of interannual environmental variability. If this intrinsic pressure is not modeled explicitly, climate anomalies can be spuriously credited for patterns that are, in fact, demographic and historical in origin. Isolating ecological momentum is, therefore, a prerequisite for determining whether warming acts primarily as a trigger for colonization or as a facilitator of expansion after establishment.

Other factors beyond water temperature and ecological momentum may positively contribute to the expansion of tropical fishes. Species originating from the Red Sea are inherently exposed to saltier water conditions compared to Mediterranean species [1]. These salinity differences suggest that recent salinification trends observed in the Mediterranean [30] may facilitate the expansion of Red Sea species by creating an increasingly suitable osmotic environment [31]. Furthermore, the spatial spread of Lessepsian fishes is non-random and proposed to be influenced by advective ocean currents. Occurrence patterns tend to increase downstream along the cyclonic coastal current regime of the Mediterranean [32,33]. For instance, the eastern Levantine Basin accumulates far more Lessepsian fish occurrences than the northwestern tip of the Egyptian coast, although both are equidistant (∼500 km) from the mouth of the Suez Canal (Fig. 1A; [34]). This suggested current-driven dispersal intensifies in the Levantine Basin, followed by the Aegean Sea, and continuing westward [35]. Additionally, decadal reversals in these large-scale circulation patterns can dynamically modulate inter-basin connectivity, potentially creating discrete windows of opportunity for species spread [36]. Finally, the physical enlargement of the Suez Canal itself can amplify propagule pressure into the region [37]; successive widening projects over the past decades have increased the Canal’s cross-sectional area, enhancing its capacity as an invasion corridor [38].

**Figure 1.**
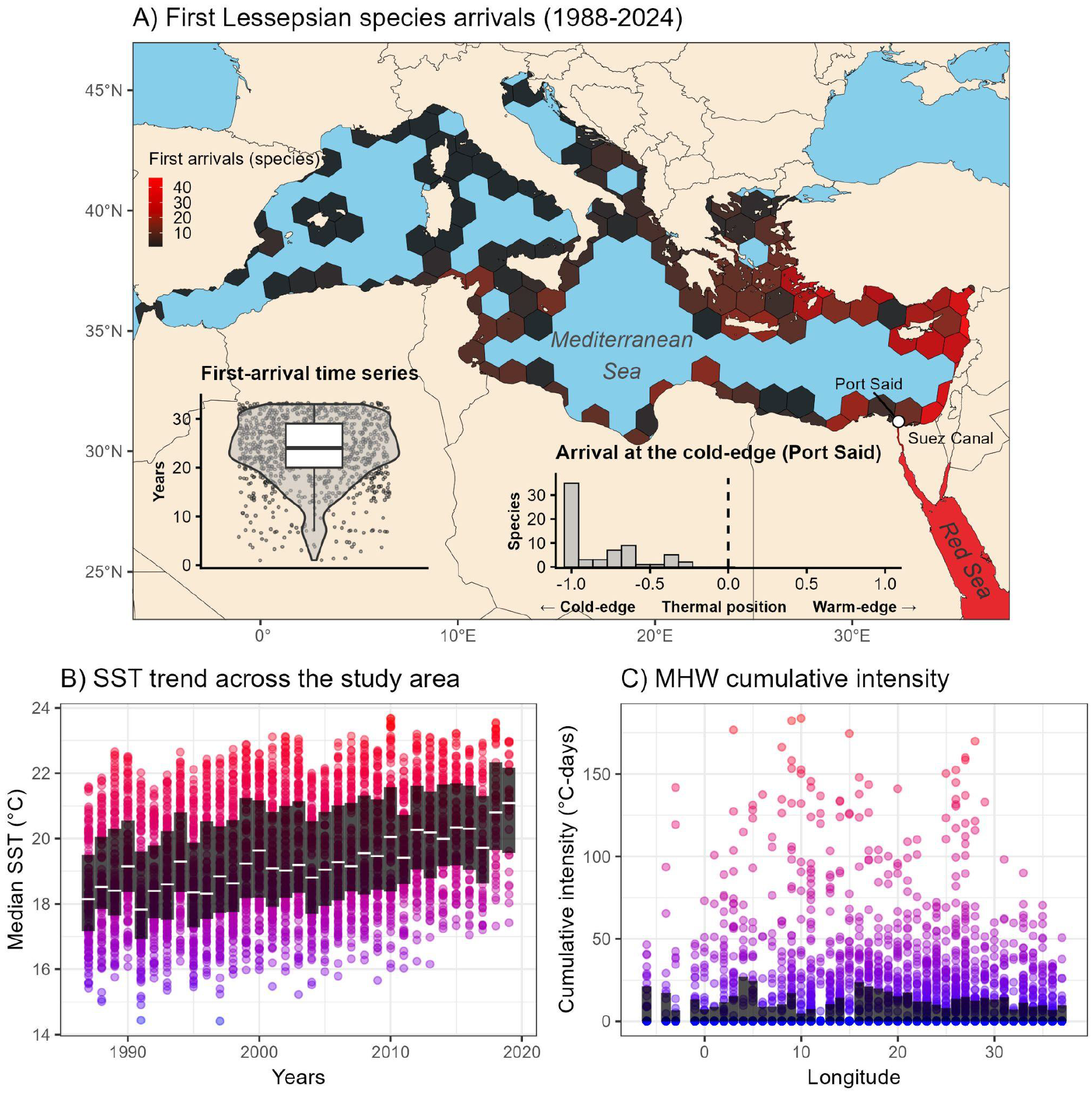
First unique species-hexagon records of Lessepsian Red Sea fish in the Mediterranean (A), long-term SST trends (B), and MHW cumulative intensity (C). A) First record count within the period of 1988-2024 (species = 75, hexagons = 134). Colors represent first-arrival count and the Red Sea is colored in red to highlight the source of Lessepsian migration. White point denotes the mouth of the Suez Canal (Port Said). The left inset boxplot denotes the distribution of the first-arrival time series (n = 1,059). Each dot represents a species within a hexagon, associating annual climatic data with records. The right inset histogram denotes the thermal position of Lessepsian species once positioned in the Mediterranean Sea at Port Said—All species are positioned within their cold edge upon entrance. B) Temporal SST trend along the recorded sites in panel A. Each year is represented by all 134 hexagons, and dots represent the median SST. Warmer colors represent warmer temperatures. Black shading represents the annual interquartile range, with white lines denoting the annual medians. Long-term warming of the entire study area is demonstrated. C) Annual MHW cumulative intensity across longitudes. Each dot represents an annual measure within a hexagon, with warmer colors denoting stronger events. Black shading represents the annual interquartile range. MHWs are spread across the western, central, and eastern Mediterranean Sea.

Taken together, the Mediterranean Sea provides a unique and stringent model system to test these competing mechanisms. It is a global hotspot for biological invasions [39,40] and for warming [41–43], with basin-wide heating rates often exceeding global averages [44]. The basin also features a pronounced longitudinal thermal gradient, with warmer conditions toward the Levantine Basin in the east, adjacent to the Suez Canal. This region acts as a physiological filter: tropical Indo-Pacific species—intrinsically warm-adapted—encounter the cold edge of their thermal range upon entering the Mediterranean (Figure 1A-right panel inset, [45]). Their subsequent spread parallels the broader global pattern of species tracking shifting isotherms [46], particularly in warm oceans [47].

Building on foundational studies of potential range expansion [34,48,49], we capture the dynamics of colonization utilizing a comprehensive spatiotemporal compilation of Lessepsian fish records. We regress these records against high-resolution daily Sea-Surface Temperature (SST; [50]), localized salinity, advective currents, and the progressive Suez Canal expansion across a continuous 32-year first-arrival framework. Specifically, we model the exact timing of first arrival by tracking local sequences of non-detections (zeros) that culminate in an arrival event (one). This data structure associates the year of arrival with our time-varying climatic and physical predictors. By evaluating this temporal pacing, we compare the effect of MHW cumulative intensity against chronic SST trends, while accounting for intrinsic ecological momentum. Contrary to prevailing emphasis on extremes, we find that redistribution in this system is dominated by intrinsic ecological momentum and consistently facilitated by chronic warming, with MHWs contributing negligibly. This means that safeguarding marine ecosystems will depend less on reactive responses to discrete heatwave events and more on sustained mitigation of chronic warming that is continuously reshaping biogeographical boundaries through the Suez Canal corridor.

## Results

To decouple the climatic drivers of species redistributions from the Red Sea, we analyzed 1,059 first-arrival time series across 75 Indo-Pacific species using a time-to-event framework with Generalized Linear Mixed Models (GLMMs). Rather than assuming static spatial equilibrium, this approach allowed us to estimate the localized colonization probability of first arrivals across the Mediterranean from 1988 to 2020 (Fig. 1). While our compiled records dataset maps the invasion front up through 2024, the statistical analysis was restricted to records through 2020 to align with the temporal coverage of our standardized sampling-effort proxy. This allowed us to rigorously control for observational biases by conditioning the first-arrival time series on this localized monitoring metric (Fig. S1), alongside the year in which each species first arrived (see ‘Accounting for monitoring effort and non-indigenous species spread pattern’ section). Additionally, to account for potential lagged biological responses to MHWs [24], climatic data were extracted starting a year prior (1987). Within this framework, we compared thermal extremes (MHW annual cumulative intensity) against chronic thermal change (annual SST). To isolate these climatic drivers, we controlled for non-climatic forces: annual sea surface salinity (SSS), the multidecadal enlargement of the Suez Canal, and advective ocean currents (Current Propagation Index; CPI). This framework allowed us to separate external environmental forcings from ecological momentum—defined here as the intrinsic demographic inertia derived from time-dependent spatial spread of populations (i.e., years). Ultimately, this enabled us to quantify whether the > 150-year Lessepsian migration is currently paced by short-term MHWs or sustained background warming.

### Ecological momentum and chronic warming govern colonization

The redistribution of Indo-Pacific fauna into the Mediterranean is overwhelmingly associated with ecological momentum and sustained ocean warming, rather than acute climatic anomalies (Fig. 2). These colonization contributors were revealed by simultaneously accounting for time, annual SST, long-term climatic baselines (SST - climate), and MHWs alongside monitoring effort, SSS, currents, and year-since-arrival within a single modeling framework. Our global model explained a substantial portion of colonization variance (i.e., the variance explained; R²m = 0.48; R²c = 0.53; Table S1), with ecological momentum (years) acting as a dominant force. The temporal component (years; i.e., representing the cumulative ecological momentum of the invasion) accounted for 65.93% of the explainable variance. This indicates that the demographic pressure associated with the Suez Canal creates a self-perpetuating invasion front that advances independently of short-term environmental variability. Among the climatic variables, annual median SST was the primary facilitator, contributing 26.65% of the explainable variance. While a region’s static historical climate (SST - climate; representing whether a spatial location is natively warmer or cooler) did not significantly determine colonization, progressive interannual warming (SST - annual) within those regions did. This is consistent with a rising thermal baseline progressively eroding physiological barriers and permitting expansion of tropical species into cooler, previously restricted latitudes and longitudes (Fig. S2).

**Figure 2.**
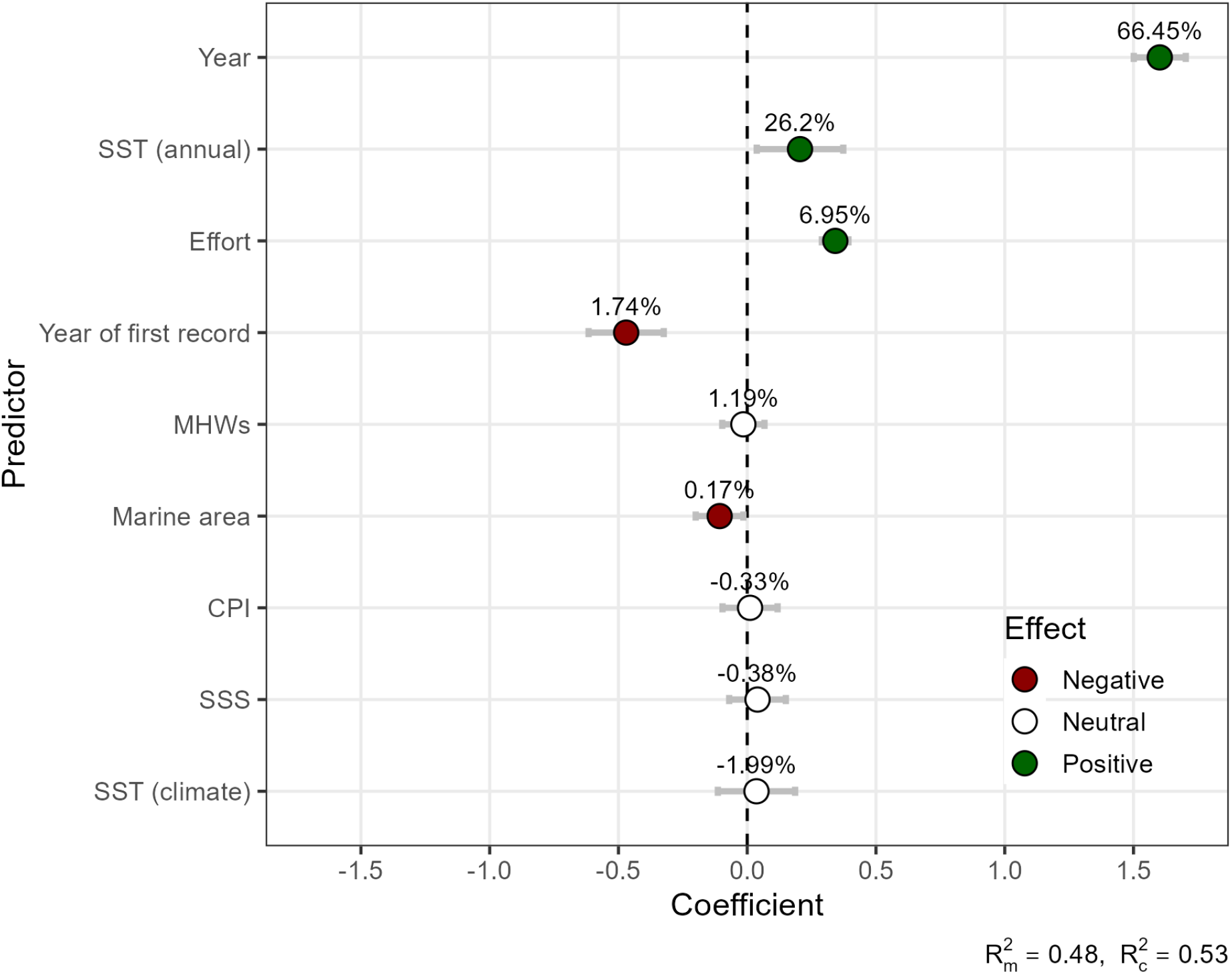
The relationship between the probability of non-indigenous Red Sea species colonization and the studied predictors (n = 24,707 presence and absence records). Points represent GLMM scaled coefficients, and error bars represent 95% confidence intervals. Predictors are ordered by the relative explained variance component of each predictor from the overall marginal variance. The vertical dashed line denotes no relationship with colonization probability. Years and annual SST account for the majority of the explained variance and positively contribute to the colonization probability. MHWs are not significantly related to the probability of observing non-indigenous colonizations.

### MHWs exert no detectable influence on colonization

MHW cumulative intensity was not associated with the colonization of tropical fish in the Mediterranean Sea (Fig. 2, Table S1). MHW cumulative intensity explained only 1.19% of the variance and showed no significant relationship with colonization probability. This result was robust to model selection: while MHW intensity appeared only in one of the top-ranked models (Δ AICc ≤ 2), it had a negligible effect and low relative importance (Table S1-S3; sum of AIC weights [SW] = 0.29), whereas ecological momentum and chronic warming were consistently retained as strong, highly significant predictors (SW of 1 and 0.98 respectively; Table S2, S3). Physical oceanographic connectivity with the Suez Canal (CPI) also showed no significant effect (Fig. 2, SW = 0.28), suggesting that the spread of the examined fishes is not dictated by passive larval drift or current regimes, but by the biological realization of their niches. Furthermore, annual SSS demonstrated no significant relationship with invasion pacing (Fig. 2, Table S1). Variance partitioning via glmm.hp assigned infinitesimally small negative shares (all magnitudes < 2%) to several of these non-significant covariates (e.g., CPI, SST - climate, and SSS; Table S1). This is expected behavior for variables that share minor background covariance but possess near-zero independent explanatory power (i.e., functional zeros acting as weak statistical noise), further confirming their negligible role in driving colonization.

### Warmer climate acts as a continuous climatic facilitator for post-establishment expansion

By segmenting the invasion process into pre- and post-establishment phases (quantitatively defined as records 1–3 versus record 4 onwards, respectively; see ‘Statistical analyses’ section), we identified a stage-dependent mechanism: chronic warming does not drive arrival, but facilitates proliferation. During the pre-establishment phase, colonization probability was primarily related to ecological momentum (64.77%), monitoring effort (measured as annual unique sources per hexagon; 13.98%), and year of first arrival (14%), while warming played no significant role (Fig. 3A, Table S4). Once populations transitioned into the post-establishment phase, the influence of the chronological variable (Year) increased to 82.64% of the explainable variance and the long-term climatic gradient (SST - climate) became a significant positive facilitator of colonization probability (Fig. 3B, Table S5), explaining 2.02% of the variance. This indicates that while arrival is comparatively stochastic or momentum driven, subsequent expansion is deterministically fueled by a warmer climate, promoting spread into areas that were previously temperature-limited.

**Figure 3.**
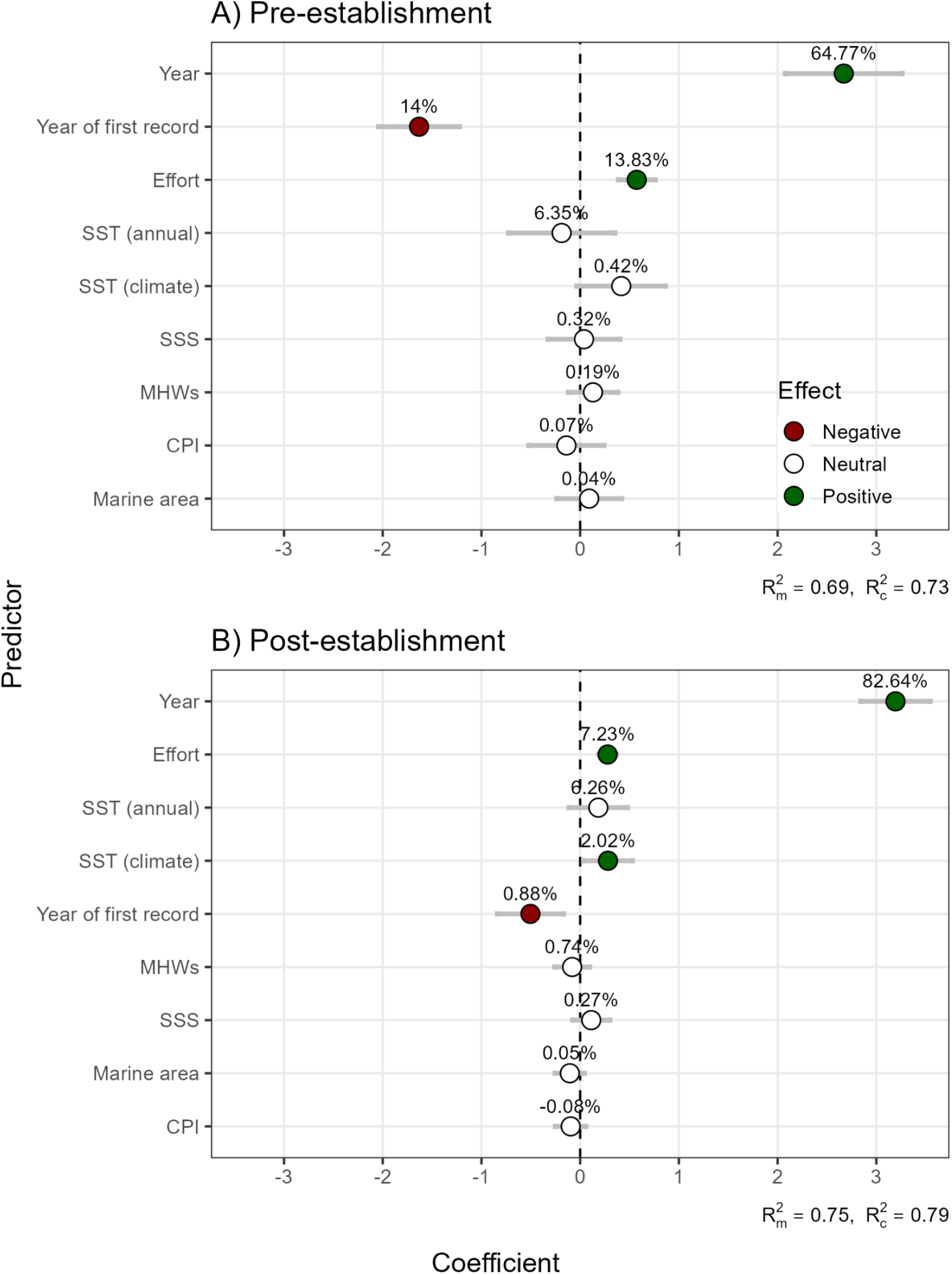
The relationship between the probability of non-indigenous Red Sea species colonization and the studied predictors across different invasion phases. Panel A represents the pre-establishment phase (n = 2,634 presence and absence records), and Panel B represents the post-establishment phase (n = 8,532). Points represent GLMM scaled coefficients, and error bars represent 95% confidence intervals. Predictors are ordered by the relative explained variance component of each predictor from the overall marginal variance. The vertical dashed line denotes no relationship with colonization probability. The temporal trend (Years) and SST (climate) show a positive effect in the post-establishment phase (B), while monitoring effort is more influential in the pre-establishment phase (A).

Finally, our results remained robust after accounting for essential non-climatic controls, confirming that the observed patterns are biological rather than observational artifacts. Monitoring effort was a significant positive predictor (6.95%), yet its influence was strongly stage-dependent: it was critical for detecting rare, pre-establishment pioneers (13.83%) and less influential once populations proliferated (7.23%). We also found a significant negative relationship with the year of first record (1.74%), indicating that old arrivals maintain higher colonization probabilities than recent arrivals. This reinforces the role of ecological momentum, consistent with time since introduction conferring a cumulative demographic advantage that operates alongside climatic facilitation. Finally, hexagons with smaller marine area—mostly representing geographically complex coastal regions (e.g., islands within the Aegean Sea)—had larger colonization probability, suggesting that Lessepsian colonization may be stronger inshore than offshore.

### Sensitivity analyses confirm the core drivers and highlight trait-mediated colonization

To ensure our conclusions were not artifacts of omitted physical or biological covariates, we performed five independent sensitivity analyses (Tables S6-10). First, explicitly controlling for spatio-temporal pseudoreplication (via a crossed hexagon-by-year random effect) confirmed that the dominance of ecological momentum (Year; p < 0.0001) and chronic warming (p = 0.025) are robust to unmeasured localized anomalies. Second, when accounting for the historical cross-sectional enlargement of the Suez Canal, the temporal effect of ‘Year’ remained the overwhelmingly dominant positive driver (p < 0.0001), confirming our chronological proxy natively captures the continuous escalation of propagule pressure. Third, we tested the previous suggestions that decadal reversals in large-scale ocean currents—specifically the Adriatic-Ionian Bimodal Oscillating System (BiOS) [36,51]—might create episodic ‘windows of opportunity’ that drive pulsed invasions. Because these circulation shifts alter how water flows between Mediterranean basins, they have been hypothesized to be a key advective driver of colonization. However, including the BiOS phase in our models did not alter our core findings: demographic momentum and chronic warming remained the dominant drivers. Interestingly, the cyclonic BiOS phase actually exhibited a highly significant negative effect on the Mediterranean-wide colonization probability (p < 0.0001). This indicates that while cyclonic current reversals might be important for pulsing certain marine organisms northward into the Adriatic, we currently find no evidence that the BiOS positively facilitates the spread of Lessepsian fishes at a Mediterranean-wide scale. Instead, for fishes, the northward deflection may temporarily interrupt the primary westward migration corridor, resulting in reduced colonization rate across the broader Mediterranean.

Finally, we evaluated species-specific thermal traits. While a species’ native thermal range (i.e., thermal generalists versus specialists) showed no significant relationship with invasion pacing (p = 0.621), the species temperature index was a significant positive predictor (p = 0.047). This indicates that while the aggregate invasion is governed by demographic momentum and chronic warming, species with warmer native thermal affinities possess a measurable colonization advantage over relatively colder-affinity migrants. This result is of particular interest as the Lessepsian species pool is intrinsically warm-adapted, nevertheless inter-specific variation in colonization response emerges.

## Discussion

The anthropogenic reconnection of the Red Sea with the Mediterranean Sea provides a large-scale case study in biotic reshuffling, functioning as a “natural laboratory” in which the rules of species redistribution can be examined under sustained propagule supply and rapid environmental change. In climate-change ecology, an increasingly influential view is that extreme warming events (e.g., MHWs) are dominant accelerants of range shifts and invasions by breaching native biotic resistance and generating short-lived windows of opportunity [9,11–13,16]. Our results challenge that framing for Mediterranean tropicalization. Across 32 years of basin-wide first-arrival time series, colonization dynamics are dominated by intrinsic ecological momentum and are consistently facilitated by chronic background warming; acute thermal extremes contribute negligibly.

A central implication is that the processes that make MHWs conspicuous—and often ecologically damaging—do not necessarily make them decisive for colonization. MHWs can coincide with mass mortality of native species [42], and, in principle, could promote invasion through a dual pathway: (i) competitive release via native species declines and (ii) transient thermal facilitation for warm-affinity taxa at their cold range edge [17]. Yet, we detect no association between MHW cumulative intensity and colonization probability. This decoupling challenge is difficult to reconcile with a model in which invasion success is primarily constrained by episodic reductions in competition [52,53]. A parsimonious interpretation is that, for many Lessepsian fishes, competitive barriers are not the limiting step because invaders frequently exploit ecological opportunities that are weakly filled by native assemblages, or occupy functional space that is effectively under-saturated [54]. Under such conditions, even substantial short-term reductions in native biomass would yield little marginal benefit for establishment or spread.

The absence of a heatwave signal also points to a more basic constraint: physiological limits and baseline thermal suitability appear to be the dominant bottleneck. Chronic warming emerged as the principal climatic facilitator, consistent with a mechanism in which the upward drift of mean conditions progressively converts marginal habitat into viable habitat, thereby relaxing cold-edge constraints over broad spatial extents. This aligns with macroecological evidence that marine taxa track shifting isotherms with higher fidelity than terrestrial taxa [4,47,55–57]. It also suggests a useful conceptual distinction between “pulse” short-lived extremes and “press” persistent baseline shifts. In this system, redistribution is governed primarily by the press component: the gradual rise of SST redefines where populations can persist and expand, whereas pulses leave little imprint on the probability of successful colonization at the spatiotemporal grain of our analysis.

Methodologically, our results help explain why the literature on heatwave-driven redistribution remains mixed. Previous foundational work by Givan et al. [58] demonstrated that thermal affinity drives local community restructuring, showing that warm-adapted species rapidly increased in abundance along the eastern Levantine coast across two discrete time periods. Building upon these insights, our study scales up to a continuous 32-year time series across the entire Mediterranean to resolve the dynamic pacing of this ongoing invasion. While prior work has utilized aggregated annual variances [59] or discrete comparisons to elucidate the spread of warm-affinity species [11,15], these approaches are distinct from capturing the temporal dynamics of range expansions. Detecting the unique contribution of extreme events requires counterfactual inference—explicit comparison against periods without MHWs under comparable background conditions [60]. In practice, many post-event studies lack long pre-event baselines or appropriate controls, and therefore risk attributing to “the event” responses that reflect ongoing secular warming. More generally, if long-term trends are not modelled explicitly, acute anomalies can be confounded with the rising mean [61]. Crucially, detrending does not mask the biological signal of MHWs if an impact indeed exists [62]; it ensures that the progressive lifting of the thermal baseline is not recorded as a continuous extreme event [61]. By analyzing continuous first-arrival time series across decades and separating detrended heatwave metrics from annual SST, we show that effects commonly ascribed to shock of extremes may, in some contexts, be dominated by the quieter influence of baseline warming.

Our stage-structured results further clarify mechanisms and resolve an apparent paradox: warming matters, but not in the way often implied by an “event trigger” narrative. The Suez Canal provides a permanent connection that sustains introduction pressure, so the first arrivals are largely governed by propagule supply and time-dependent ecological momentum. In contrast, warming exerts its strongest influence after establishment, acting as a climatic facilitator that accelerates expansion once populations have crossed initial demographic and ecological threshold. This pattern is consistent with a physiological release model in which population performance increases as recipient environments approach—yet remain below—thermal optima characteristic of source regions [63], producing predictable spatial gradients in spread potential [59]. Critically, this implies that models focused solely on extremes can miss the more deterministic component of tropicalization driven by baseline shifts and demographic inertia.

The broader biogeographical context matters. The scale of redistribution shows that anthropogenic corridors can operate as geological-scale interventions, reversing long-standing barriers and enabling inter-basin exchange [64–66]. In this context, the Mediterranean tropicalization serves as an empirical evidence of the anthropogenic “global bioflow” [67]. This system illustrates that vector-driven introductions (via the Suez Canal) and climate-driven range expansions (facilitated by warming seas) are not disparate phenomena, but rather interconnected pathways of biological invasion. We argue that the resulting invasion dynamics are not explained by a single factor— salinity contrasts [31], currents [38], preadaptation of Red Sea biota [68], or niche availability [54] but by their further interaction with sustained warming and demographic feedback. The demographic reservoir established in the Levantine Basin can generate a self-reinforcing invasion front advancing westward [35], largely independent of short-term anomalies. The lack of a detectable effect of oceanographic connectivity suggests that the spread is not simply a function of passive larval drift. Instead, post-settlement bottlenecks—recruitment, resource acquisition, habitat matching, or biotic filtering—may determine whether arrivals translate into persistent occupancy, as observed even within native ranges [69]. Indeed, climate alone does not guarantee a successful expansion into the Mediterranean [70]; establishment in this system also relies on inherent biological traits [71,72], such as a species’ capacity for post-invasion habitat lability to exploit novel non-coral environments [73]. This interpretation is consistent with the dominance of ecological momentum: once populations are established, local proliferation and stepping-stone dynamics can overwhelm variability in transport. Further, our diagnostics confirm that the pacing of this invasion is not an artifact of unmeasured monotonic forcing. By modeling salinity trends, episodic current shifts (e.g., BiOS), and canal enlargement, we demonstrate that years isolate the ecological momentum of the invasion front, separate from the shifting properties of the basin.

Several caveats delimit inference and point to future work. First, while our core conclusions represent the aggregate, macroecological signal for Lessepsian fishes, our sensitivity analyses confirm that individual responses are also mediated by species-specific biology. Specifically, while physiological plasticity (native thermal range) did not significantly modulate invasion pacing, species with warmer native thermal affinities (higher species temperature index) exhibited significantly higher colonization probabilities. This empirical finding at the Mediterranean scale directly complements localized community restructuring observed by Givan et al. [58], confirming that thermal affinity acts as an internal biological filter interacting with external climatic drivers. Other traits such as dispersal modes, reproductive life histories, and post-invasion habitat lability [73] inherently interact with these climate drivers to determine event responses. Second, our database inherently aggregates across a wide variety of unstandardized sampling methods (e.g., commercial landings, scientific trawls, and underwater visual censuses). While modeling binary first-arrivals at a large spatial grain (∼12,000 km² hexagons) mitigates severe gear-habitat biases, and our random effects structure absorbs baseline differences in species detectability, this opportunistic sampling still introduces observation noise. Third, we quantify MHWs using surface metrics; other metrics, such as subsurface extremes or event timing relative to sensitive life stages could matter even if annual cumulative intensity does not [74,75]. Fourth, colonization probability based on occurrence records may be insensitive to short-lived abundance pulses that do not translate into persistent establishment. These limitations do not weaken the central result, but they clarify where a heatwave signal could plausibly manifest: not as a primary determinant of colonization success at basin scale, but potentially as a modifier of demographic rates, impacts, or community consequences once populations are present.

Taken together, the Mediterranean-Red Sea corridor suggests a generalizable template for redistribution under global change: sustained connectivity supplies propagules, ecological momentum supplies inertia, and chronic warming supplies the directional environmental facilitation that sets the pace and geography of spread. In this view, while MHW impacts are documented worldwide [60,62,76], they are not the principal architect of tropical colonization in this system. The dominant drivers are less episodic and less visible: the persistent rise of thermal baselines, compounded by the bridges we have built between biomes.

## Methods

### Collection of geo-referenced records

Data for the geo-referenced occurrence records of non-indigenous Red Sea fish species in the Mediterranean were assembled over a period of approximately 20 years. The starting point was the first edition of the Atlas of Exotic Fishes in the Mediterranean [77], with the dataset continuously curated, based on published and grey literature, until November 2025. This curated dataset forms part of the Hellenic Centre for Marine Research (HCMR) offline database. The literature search involved a retrospective review of occurrence records by species and a continuous, systematic search of all invasion-focused and Mediterranean regional journals. This effort was complemented by non-duplicate records sourced from the Occurrence Records of Mediterranean Exotic Fishes (ORMEF) database [33]. The backbone of the final database (14,175 total records) consists of peer-reviewed publications, which together amount to approximately 60% of the records [78,79]. Specifically, 4,280 records of the 75 selected non-indigenous species are based on 680 peer-reviewed publications (see supplemental methods section ‘Species selection’). Additional secondary sources contributed 1,810 records or 12% of the total. These secondary sources include the ELNAIS database [80], Global Biodiversity Information Facility (Table S6) [81] and OBIS databases [82,83], research-grade observations in iNaturalist, MEDITS surveys, the HCMR Fish Collection system, the Mediterranean Marine Invasive Species platform, and data shared by non-governmental organizations. All duplicate records were checked and removed before inclusion.

A single occurrence record was entered for each unique combination of species, spatial coordinates, and point in time. For instance, a single multi-species survey reported in one reference could yield multiple occurrence records for several species. The temporal resolution used in this work is the year of collection as documented; if the collection year was not provided, the publication year of the record was used instead. Similarly, when exact coordinates were not provided in the original publication, we manually approximated the location using Google Earth based on available contextual metadata (e.g., specific bay names, depth contours, distance from the coast, or by georeferencing published maps). These approximations were rare and were primarily limited to the oldest first record publications originating from the southeastern Mediterranean Sea. As our spatial standardization procedure bins all coordinates into a coarse hexagonal grid (averaging 12,393 km² per cell), these fine-scale manual approximations fall within the correct overarching spatial unit, preventing any impact on the spatial accuracy of the model. Accidental species misspellings were harmonized using the ‘bdc_query_names_taxadb’ function of the ‘bdc’ R package [84].

### Accounting for monitoring effort and non-indigenous species spread pattern

Analyzing raw record occurrence data may be prone to biases, including uneven spatial reporting and localized oversampling [85]. To spatially standardise occurrences and mitigate these geographic biases across the Mediterranean, we binned the raw occurrence point coordinates into Discrete Global Grid System. Specifically, we utilized Uber’s H3 Hierarchical spatial index at resolution 3 [86], which patterned the study area into comparable hexagonal cells (average area of 12,393 km^2^). Here, we maintained the year of the first species record within a hexagon and within the temporal range of the available climatic data, ranging from 1987 to 2023. To account for the varying extent of available marine area across highly fragmented coastal margins versus open-ocean environments, we intersected the hexagonal grid with global landmass polygons to extract the effective marine area for each hexagon (km²; 1st quartile = 5,131, median = 9,996, 3rd quartile = 12,582). This exact area was subsequently log-transformed and scaled for use as a covariate in our models. To further account for unequal monitoring effort, we produced a proxy variable based on the ORMEF dataset. Here, we filter hexagons represented in our record data and ORMEF, and count the unique sources (e.g., publications) reporting new non-indigenous species occurrences within each hexagon and year (Fig. S1; including absence of sources). This monitoring effort proxy variable (integer) was added to our models as a predictor to separate the signal of true non-indigenous species spread from that of increased monitoring effort (see ‘Statistical analyses’ section). In addition, as older arrivals are predicted to have more occurrences than more recent arrivals, the year of the first record was recorded and incorporated as a predictor in the analyses.

To account for advective cyclonic dispersal of Lessepsian fishes, we developed a Current Propagation Index (CPI) using annual surface velocity data from the Copernicus Marine Service dating from 1987 to 2023 [50]. This spatial index was incorporated into our models to isolate the effect of advective currents from other ecological trends (see ‘Statistical analyses’ section). The CPI was calculated for every hexagon as CPI = A × B × C, where A represents inverse path distance, B represents propagated current alignment, and C represents exponential distance decay. Surface current velocity estimates were extracted at a horizontal grid resolution of 1/24° [50]. These estimates provide the Eastward and Northward current velocity components for each hexagonal cell. The inverse path distance component (A) was calculated as 1/(*d*+0.001), where *d* represents the minimum number of hexagon steps from the Suez Canal. The propagated current alignment component (B) introduces a chain-reaction mechanism where each hexagon’s alignment is influenced by adjacent hexagons closer to the source—the mouth of the Suez Canal. First, local current assistance is quantified for each hexagon by calculating the dot product between its prevailing ocean current vector and the ideal straight-line dispersal vector originating from the Suez Canal (where positive values indicate assisting flow and negative values indicate hindering flow). Second, to capture continuous propagation, the algorithm evaluates hexagons in order of distance from the Canal. A target hexagon’s final alignment score is calculated as a weighted average of the current alignment of its immediate ‘upstream’ neighbors (70%) and its own local current alignment (30%).

This algorithm rewards regions downstream of the canal (e.g., the Levantine coast) where currents consistently chain together to facilitate dispersal, while penalizing regions where currents oppose spread. The distance decay component (C) was calculated as e^(-λ×A)^. This penalty ensures that the index weighs against long-distance dispersal, reflecting the inherent ecological resistance of colonization over great distances. Consequently, a positive CPI value indicates a hexagon is supported by current propagation and close to the source, while a negative CPI value indicates strong opposition from the current, typically observed in westward locations against the prevailing counter-clockwise circulation (Fig. S3).

### Climatic variables

To estimate the effect of long-term SST trends and MHWs on the probability of Lessepsian species colonization, we extracted daily SST data from the Mediterranean Sea Physics Reanalysis at a horizontal grid resolution of 1/24° dating from 1987 to 2023 [50]. Data were extracted for all hexagons containing record data (Fig. 1A) using the ‘cms_download_subset’ function of the ‘CopernicusMarine’ R package [87]. Because mapping a square raster grid (1/24°) onto a hexagonal grid creates boundary overlaps, we utilized a spatial intersection to prevent data from “leaking” across adjacent hexagons. Specifically, the center coordinates of every 1/24° Copernicus grid cell were assigned to a specific H3 hexagon using the ‘h3jsr’ R package [88]. Grid cells whose center points fell outside the boundaries of a target hexagon were discarded. The daily SST values of all retained grid cells were then spatially averaged to produce a single daily SST value per hexagon. Finally, to reduce the increased daily temperature variability of the uppermost surface layer available, surface temperature was extracted and averaged for the first 10 meters of the water column. Similarly, to account for the local osmotic environment encountered by Red Sea migrants, we extracted monthly Sea Surface Salinity (SSS; measured in Practical Salinity Units) from the same Copernicus Reanalysis product. We applied the identical spatial intersection procedure and 10-meter depth integration to maintain precise spatial and environmental consistency. These data were then aggregated to calculate the annual median salinity per hexagon, serving as a continuous predictor of osmotic suitability in our models (Fig. S4).

To account for the impact of the background long-term SST trend, we extracted the median annual SST per hexagon and used it as a predictor in our models (Fig. 1B). To visualize SST warming rates across hexagons, we used linear models with mean SST as the response and years as the predictor (Fig. S5A). As non-indigenous species invade through the Suez Canal towards the southeastern Mediterranean, they initially settle in warmer regions. To better disentangle the effect of long-term SST trends (SST-annual) and the spatial SST gradients across the Mediterranean Sea (SST-climate), we also extracted the median maximum SST per hexagon over the years 1988–2023 (Fig. S5B). We used the median maximum SST due to its low Variance Inflation Factor value in our models (see ‘Statistical analyses’ for more details, VIF = 2.37).

To estimate the annual MHWs’ cumulative intensity (°C-days), we used a fixed reference period across 36 years, ranging from 01/01/1987 to 31/05/2023. We chose cumulative intensity in favor of other impact metrics (e.g., intensity, duration) because it integrates both the temporal and magnitude components of exposure, which are crucial for associating species impact by potential thermal extremes [89]. MHWs were defined in cases where the anomaly exceeded the 95^th^ percentile threshold and lasted more than five consecutive days. As we used the 95^th^ percentile, even a relatively weak event may represent a relatively strong MHW, and further enables comparison across other fish-related MHW-impact studies [24,62]. The daily SST data were linearly detrended prior to MHW detection to remove long-term SST trends, thus focusing primarily on SST anomalies and removing the effect of longer and more frequent MHWs throughout the end of a climatic time series [61]. This linear detrending is necessary to isolate the amplitude of short-term extreme events from the background decadal warming trend. By neutralizing the shifting mean, we ensure the model does not mix the effects of sustained, gradual warming with acute thermal extremes, allowing for a strict causal disentanglement between the two competing ecological drivers [60,61]. MHWs’ cumulative intensity was estimated using the ‘heatwaveR’ R package. Here, cumulative intensity was summed within years and hexagons, thus representing local events (Fig. 1C, Fig. S5C).

### Lessepsian species thermal position at entry

To quantitatively test the assumption that Lessepsian migrants face a physiological thermal restriction upon entering the Mediterranean, we estimated each species’ thermal position at the Suez Canal exit (Port Said) relative to its native global species temperature index [90]. To reconstruct native thermal preferences independently of the invaded range, we downloaded global occurrence records for all 75 modeled species from the Global Biodiversity Information Facility (GBIF; [91]) using taxonomic backbone matching. To ensure high data fidelity, occurrences were subjected to a rigorous automated cleaning pipeline (Table S11). All occurrences within the Mediterranean and Black Seas were excluded using a custom spatial polygon terminating precisely at the northern entrance of the Suez Canal. The remaining occurrences were filtered for marine-only environments, coordinate uncertainty (≤ 50 km), and spatial outliers, followed by spatial thinning at a 0.1° grid resolution to mitigate localized sampling bias. The final dataset included sufficient non-Mediterranean occurrence records for 66 out of the 75 modeled species.

To quantify the species temperature index, we extracted the annual global climatic SST available (1993–2016 reference period, 0.083° resolution) from the Copernicus Marine Service (GLORYS reanalysis; [92]) for every cleaned native occurrence. For each species, we calculated the minimum, median, and maximum native SST, thermal range, as well as the thermal mid-point. To determine where the Mediterranean entry environment sits relative to each species’ physiological capacity, we extracted the mean climatic SST at Port Said (using a 10 km spatial buffer). The thermal position index at entry was calculated as the difference between the local Port Said SST and the species’ median native SST, normalized by the upper half of its thermal range (cf., [62]). This index was subsequently winsorized to bounds of −1 and +1, where −1 indicates the species is experiencing its absolute cold-edge limit upon entry, 0 represents its thermal optimum, and +1 indicates its absolute warm-edge limit (Fig. 1A - right inset).

### Statistical analyses

To estimate the effect of long-term Mediterranean Sea warming and MHWs on non-indigenous species colonization probability, we employed a GLMM using the ‘lme4’ R package [93]. Specifically, we analyzed 1,059 first-arrival time series for 75 non-indigenous fish species (Fig. 1A - left inset panel). We define colonization probability as the conditional probability of a species being recorded in a hexagon for the first time, given its prior absence. The response variable, records, representing the first occurrence (1) or the absence (0) of a Lessepsian species observation within a given hexagon, was modeled using the binomial distribution with a logit link function. The data were structured such that every row represented a unique hexagon-species-year combination. For any given species within a specific hexagon, the temporal sequence consisted of pre-arrival non-detections (0) culminating in the year of first observed arrival (1), after which the sequence for that specific species-hexagon combination terminated. The fixed effects used were categorized into three groups: (1) Climatic predictors focused on acute and long-term temperature change, including detrended MHW cumulative intensity, annual median SST (SST-annual), and the long-term median maximum SST (SST-climate) estimated at the hexagon level. To account for lagged responses to acute thermal stress, MHW cumulative intensity was temporally offset by one year, such that colonization probability in year *t* was regressed against the MHW cumulative intensity of year *t-1* [24,62]. Dynamic interannual predictors (e.g., annual median SST, SSS) were evaluated for the concurrent year *t* to reflect immediate environmental suitability, while the long-term climatic baseline (SST-climate) was applied as a static spatial constant for each hexagon. (2) Spatio-physical control included the *CPI* to isolate the effect of current-assisted dispersal and geographical proximity from the Suez Canal mouth to the investigated target hexagon. (3) Ecological and monitoring control variables included the years, year of first Mediterranean record for the species, and the annual sum of sampling sources per hexagon (i.e., proxy for monitoring effort). Years (continuous) were used to estimate the ecological momentum; the year of first Mediterranean record (i.e., since the opening of the Suez Canal) was used to account for the effect of new arrivals compared to old arrivals.

We included random intercepts for hexagons and species. The hexagon random intercept accounted for spatial autocorrelation and unmeasured site-specific factors associated with the records. The species random intercept accounted for inherent, unmeasured species-level differences in colonization success and baseline detectability (e.g., absorbing the variance between highly detectable commercial species versus cryptic taxa). All continuous predictors were scaled before analysis by subtracting the mean and dividing by the standard deviation. To manage the trade-off between model fit and complexity, we used model selection with second-order Akaike information criterion [94]; AICc). We applied the ‘dredge’ function from the ‘MuMIn’ R package, to find the most parsimonious set of predictors [95]. The relative importance of each predictor was assessed using the summed AIC weights using the ‘sw’ function of the ‘MuMIn’ R package.

To estimate whether the predictors’ effect on Lessepsian species colonization changes for different phases of introduction, we tested two additional models having the same settings but different subsets. First, we trimmed the dataset to only include species that entered the Mediterranean Sea for the first time after the earliest climatic data available (i.e., after 1987). Next, we divided the data based on the records for each species into two distinct phases: (1) pre-establishment phase, comprising records for a species ranging from the first Mediterranean record up to and including the third successive chronological record, representing the initial stages of introduction. (2) post-establishment phase, comprising records from the fourth chronological occurrence onwards. This threshold builds upon the criteria established by ref [96], ensuring that the post-establishment analysis focuses on fish populations that have already demonstrated the capacity for persistence and are actively expanding.

For all models, the relative contribution of each fixed effect to the overall R^2^m was estimated using the glmm.hp function of the ‘glmm.hp’ R package [97]. This algorithm decomposes the total marginal R*^2^* into the unique and average shared variance contributed by each fixed effect across all possible sub-models. Because our model utilizes a binomial error distribution with a logit link, this decomposition strictly applies to the latent (link) scale, where fixed effects are linear and additive. While this provides a robust relative ranking of predictor importance, we caution that variance components partitioned on the latent scale cannot be directly translated into absolute additive probabilities on the observational scale. We consider this acceptable for our framework, as our objective relies on identifying the comparative dominance of background warming over episodic extremes, a relative ranking that remains robust on the latent scale. The marginal and conditional R^2^ were calculated following the framework of Nakagawa and Schielzeth [98]. The distribution-specific residual variance on the latent scale was approximated using the theoretical method [98]. Finally, overall model validity was assessed using the “simulateResiduals” function of the ‘DHARMa’ R package ([99]; Fig. S6). Multicollinearity was assessed using VIF analysis where values consistently remained below a highly conservative threshold of three; Table S13).

### Sensitivity analyses

To test the robustness of our findings against additional physical and biological covariates, we performed five independent sensitivity GLMMs while preserving the set predictors of the core abovementioned model. First, to ensure our temporal variable (Year) did not spuriously absorb the variance of co-trending physical forcing, we tested the enlargement of the Suez Canal (cross-sectional area) to account for increasing propagule pressure. Second, to ensure our conclusions regarding episodic forcing were not limited to thermal extremes, we conducted a sensitivity analysis incorporating episodic advective anomalies. The BiOS of the Ionian Sea undergoes decadal reversals between cyclonic and anticyclonic phases [51]. We integrated a two-level categorical BiOS-phase indicator into our global GLMM. To strictly align with the high-confidence satellite altimetry era from which these reversals are derived [100], this model was restricted to the 1993–2020 temporal window. Third, to control for species-specific thermal affinities, we integrated the species temperature index (see “Lessepsian species thermal position at entry” section), distinguishing between warm- and cold-water affinities following the framework of Givan et al. [58]. Fourth, to evaluate whether physiological plasticity modulates invasion pacing, we included each species’ native thermal range, defined as the difference between the maximum and minimum temperatures extracted from their global, non-Mediterranean occurrences. Finally, to rigorously account for potential spatio-temporal pseudoreplication—where unmeasured, transient local anomalies or sampling events might simultaneously affect multiple species in a given location and year—we incorporated a crossed random effect for the interaction between spatial location and time (hexagon by year). Each of these five modifications was added independently to the core binomial GLMM to verify that the primary effects of demographic momentum and chronic warming remained consistent despite these additional controls.

## Supplemental Material

### Supplemental Figures

**Figure S1.**
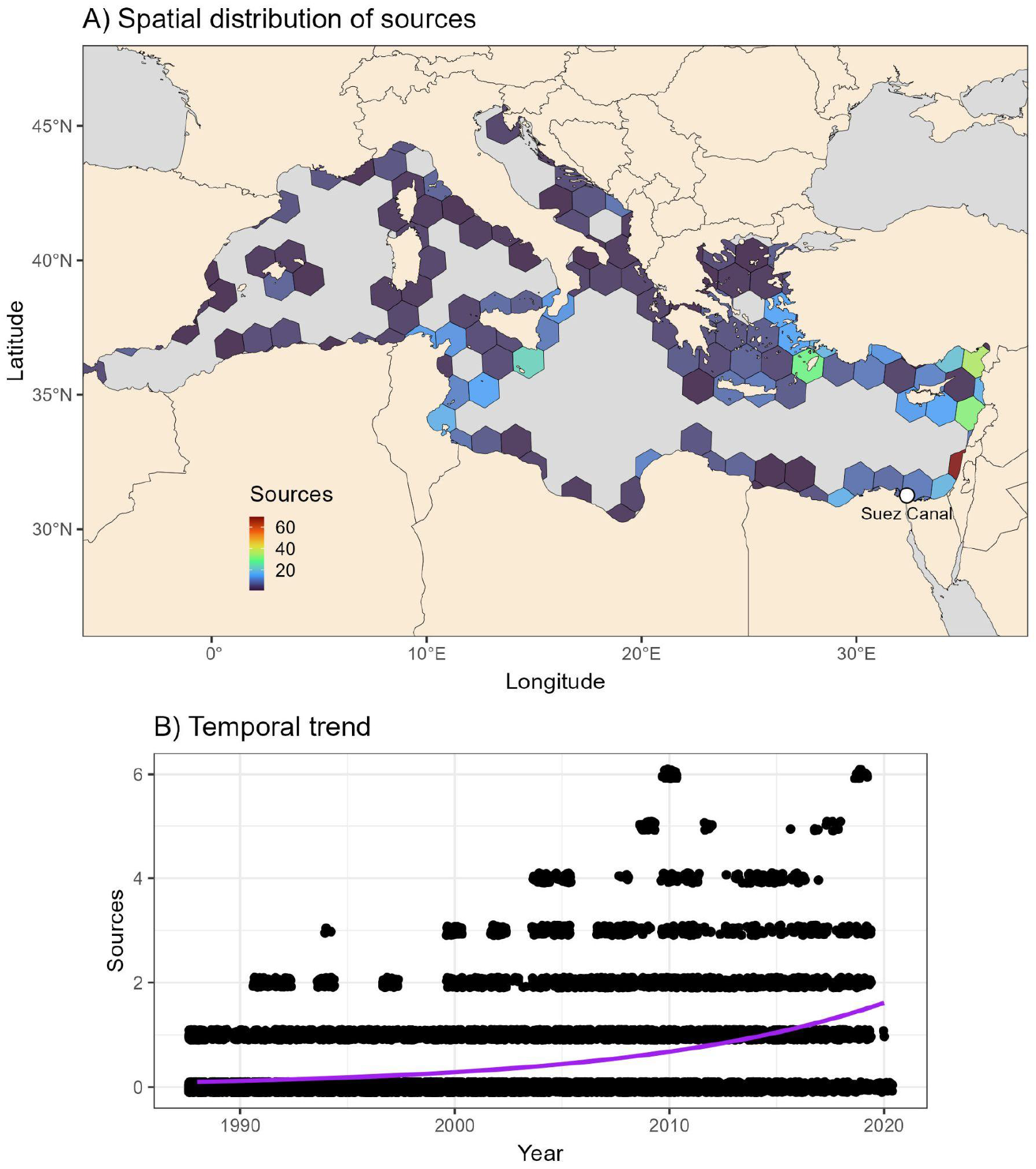
Monitoring effort proxy. A) The distribution of non-indigenous species record sources aggregated across years. The Eastern Mediterranean Basin shows more sources compared with the Western Mediterranean Basin. White point denotes the mouth of the Suez Canal. B) The relationship between summed unique sources within hexagons against years, showing increased source count over time. Note that points are jittered to better represent the data.

**Figure S2.**
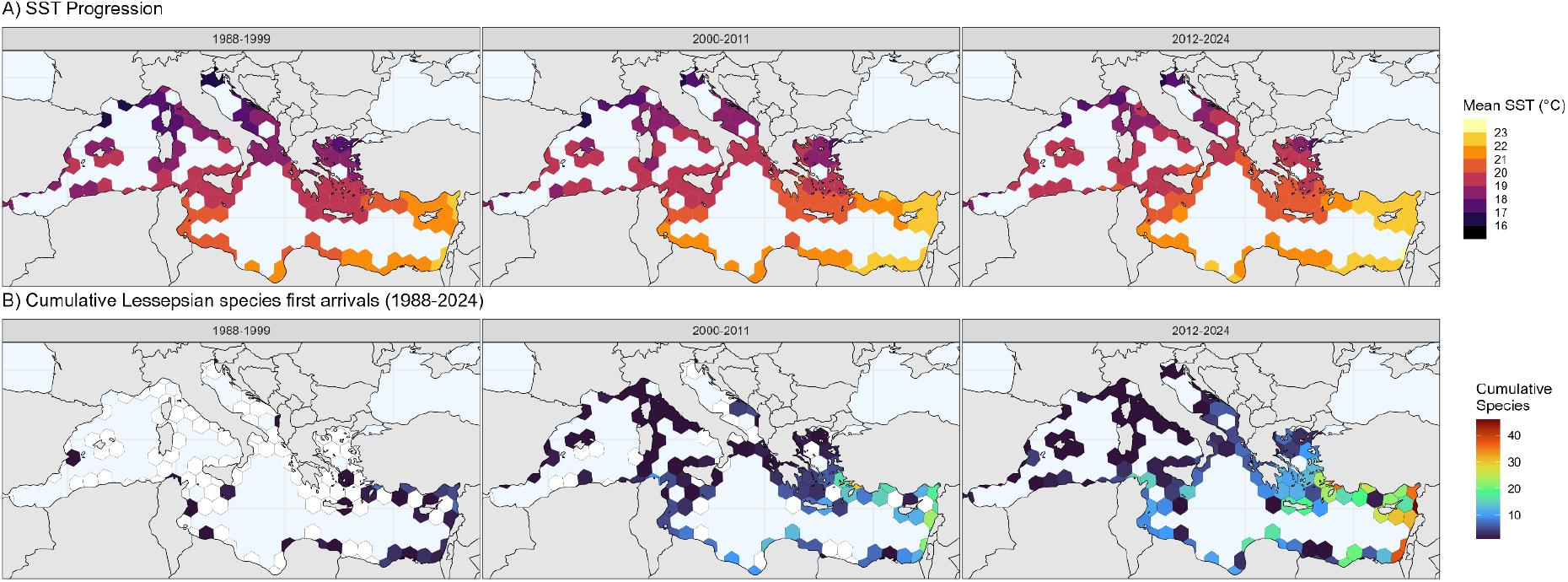
Spatio-temporal progression of SST and cumulative Lessepsian fish expansion across the Mediterranean Sea (1988–2024). (A) Decadal shifts in mean SST (°C) aggregated within the H3 hexagonal spatial grid. Temperatures are binned into discrete isotherms to visually highlight the progressive westward and northward expansion of the warming thermal envelope across three distinct time periods restricted to the temporal window of the ORMEF dataset (1988–1999, 2000–2011, and 2012–2024). (B) The corresponding cumulative species richness of first-arrival events for the 75 modeled Lessepsian fishes across the same three temporal periods. Hexagons with zero recorded species are masked (white) to emphasize the active invasion front. These panels demonstrate a synchronized, basin-wide spread of the tropical species pool that closely tracks the shifting chronic climatic baseline.

**Figure S3.**
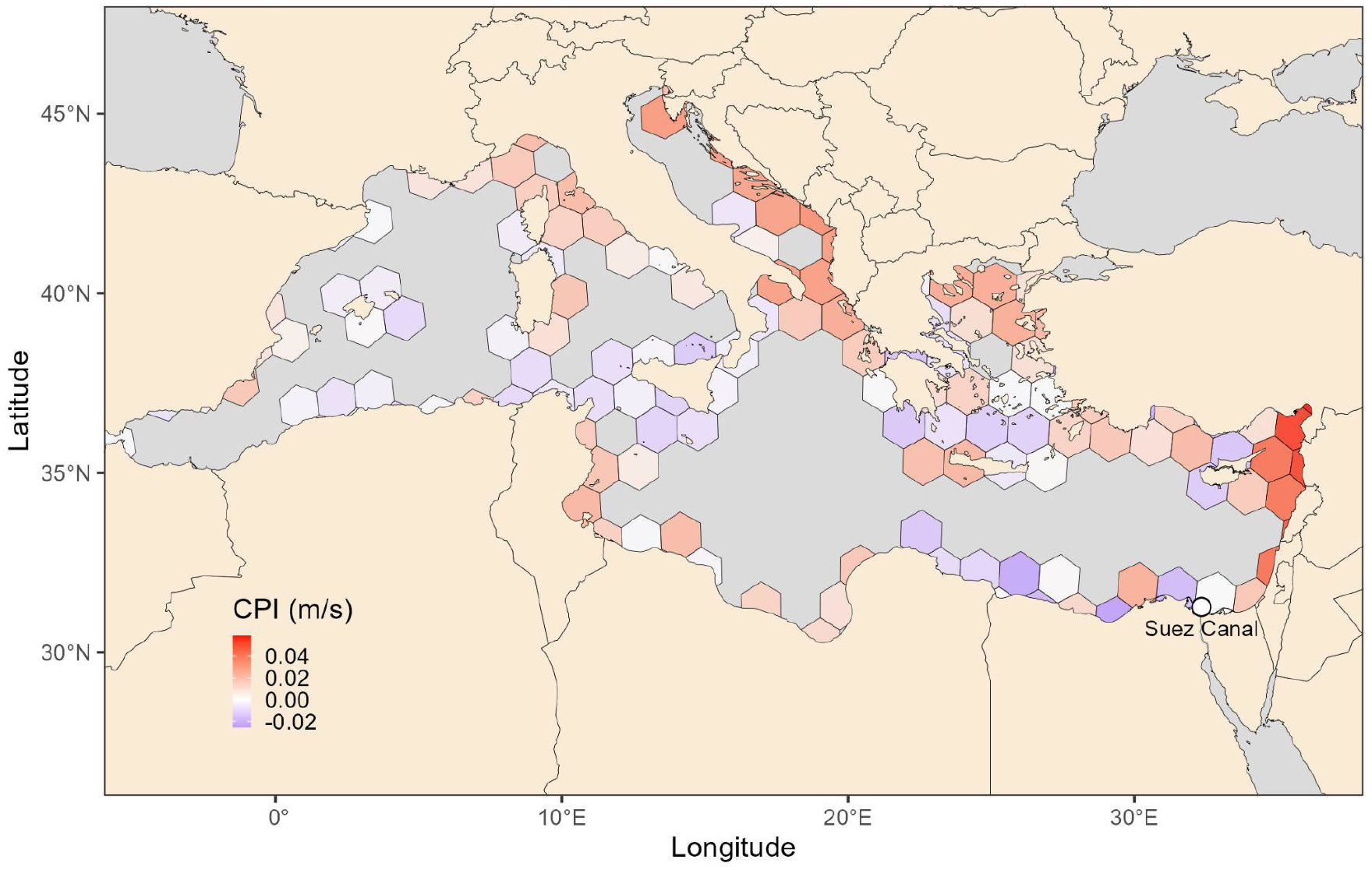
Current Propagation Index (CPI) represents the integrated dispersal potential from the Suez Canal source for each Mediterranean hexagon. Positive CPI values (warm colors) indicate that a hexagon is supported by current-assisted propagation and proximity to the mouth of the Suez Canal (i.e., white point) through successive hexagons. Negative CPI values (cold colors) indicate strong opposition from currents, typically observed in westward locations against the prevailing circulation patterns. The CPI incorporates chain-reaction propagation where hexagons influence their downstream neighbors, creating persistent dispersal corridors or barriers. White point denotes the mouth of the Suez Canal.

**Figure S4.**
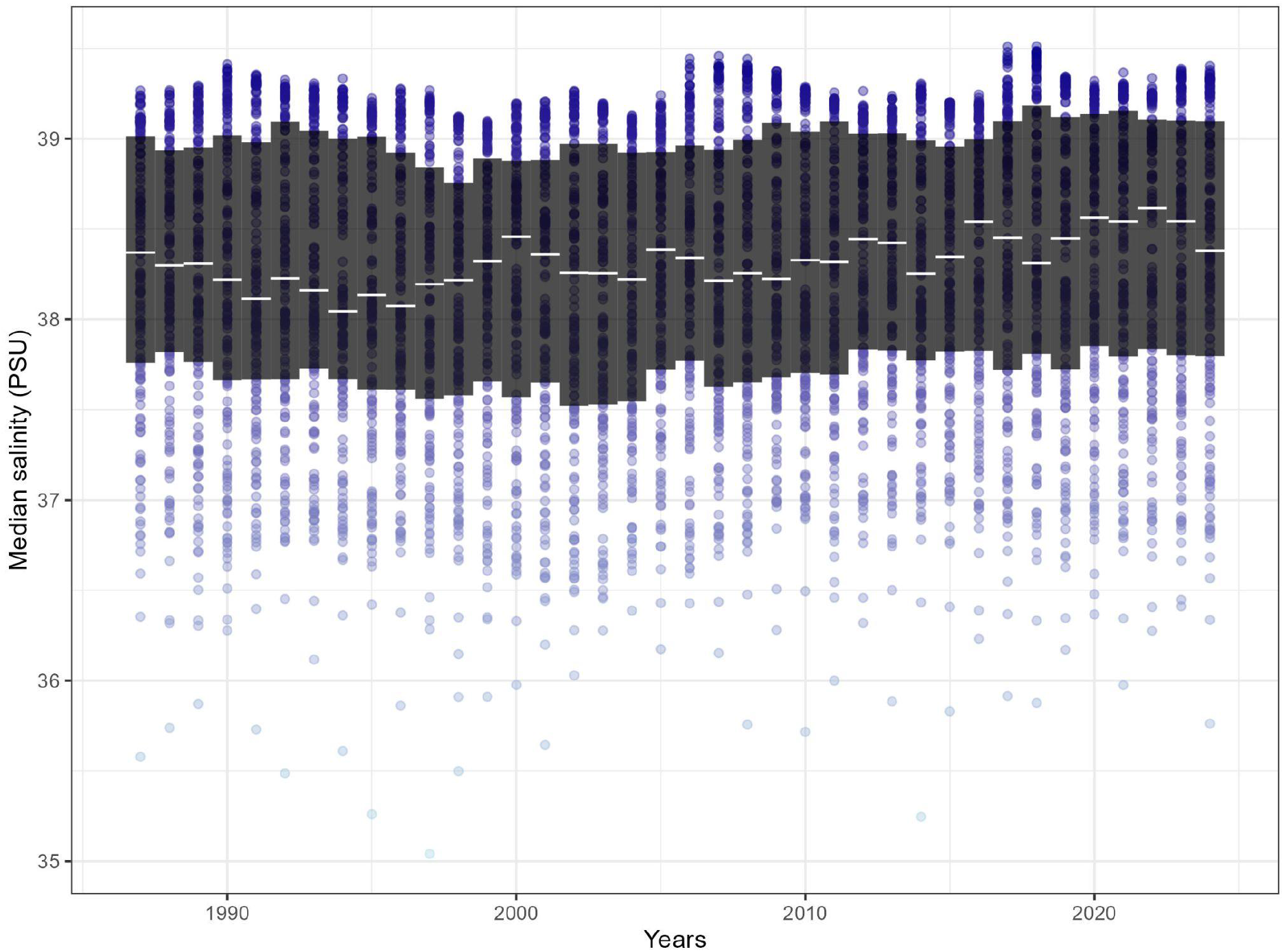
Spatiotemporal trend of annual surface salinity across the Mediterranean study area. To account for progressive salinification, Sea Surface Salinity (SSS, measured in PSU) was extracted from the Copernicus Marine Service Mediterranean Sea Physical Reanalysis product (MEDSEA_MULTIYEAR_PHY_006_004). Data were derived from the monthly 4.2 km resolution layer (cmems_mod_med_phy-sal_my_4.2km_P1M-m) and integrated across the upper mixed layer (0–10m depth). The plot illustrates the distribution of the calculated annual median surface salinity for every specific hexagon across the 36-year study period. Background points and colored gradients reflect the local salinity of individual hexagons, while the black boxplots and white crossbars denote the spatial variance and overall annual median across the basin for a given year. This highly resolved, hexagon-specific continuous proxy was subsequently integrated into the GLMMs to test the local osmotic suitability hypothesis against intrinsic ecological momentum, MHWs’ cumulative intensity, and chronic warming.

**Figure S5.**
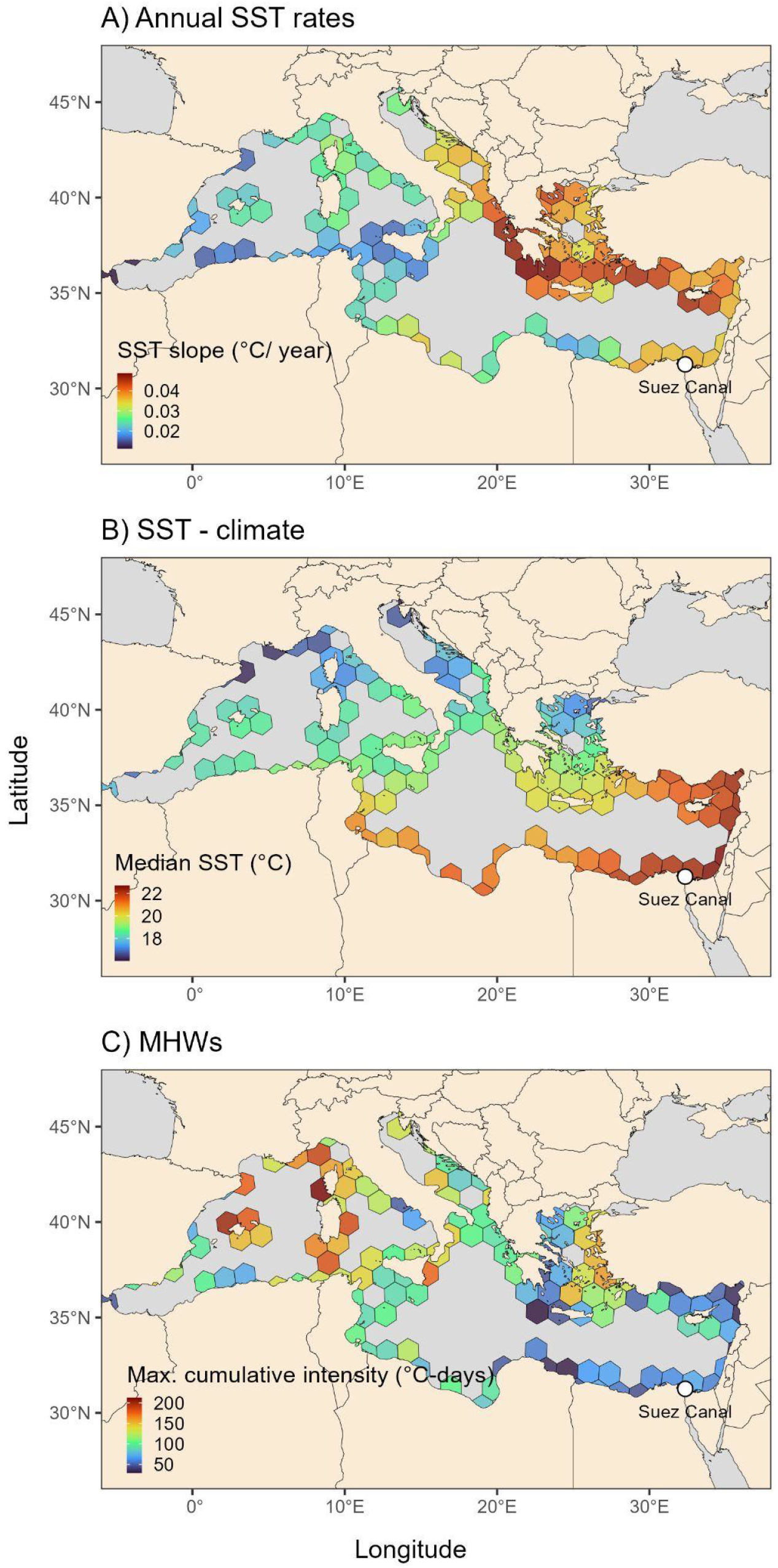
Spatially-summarised SST warming rates (A), median SST (B), and maximum cumulative intensity of detrended MHWs (C). Data is summarised within each hexagon over the years 1987-2023 with warm colors representing higher warming rates, median SST, or high maximum cumulative intensity. Cold colors represent lower warming rates, median SST, and maximum cumulative intensity. White point denotes the mouth of the Suez Canal.

**Figure S6.**
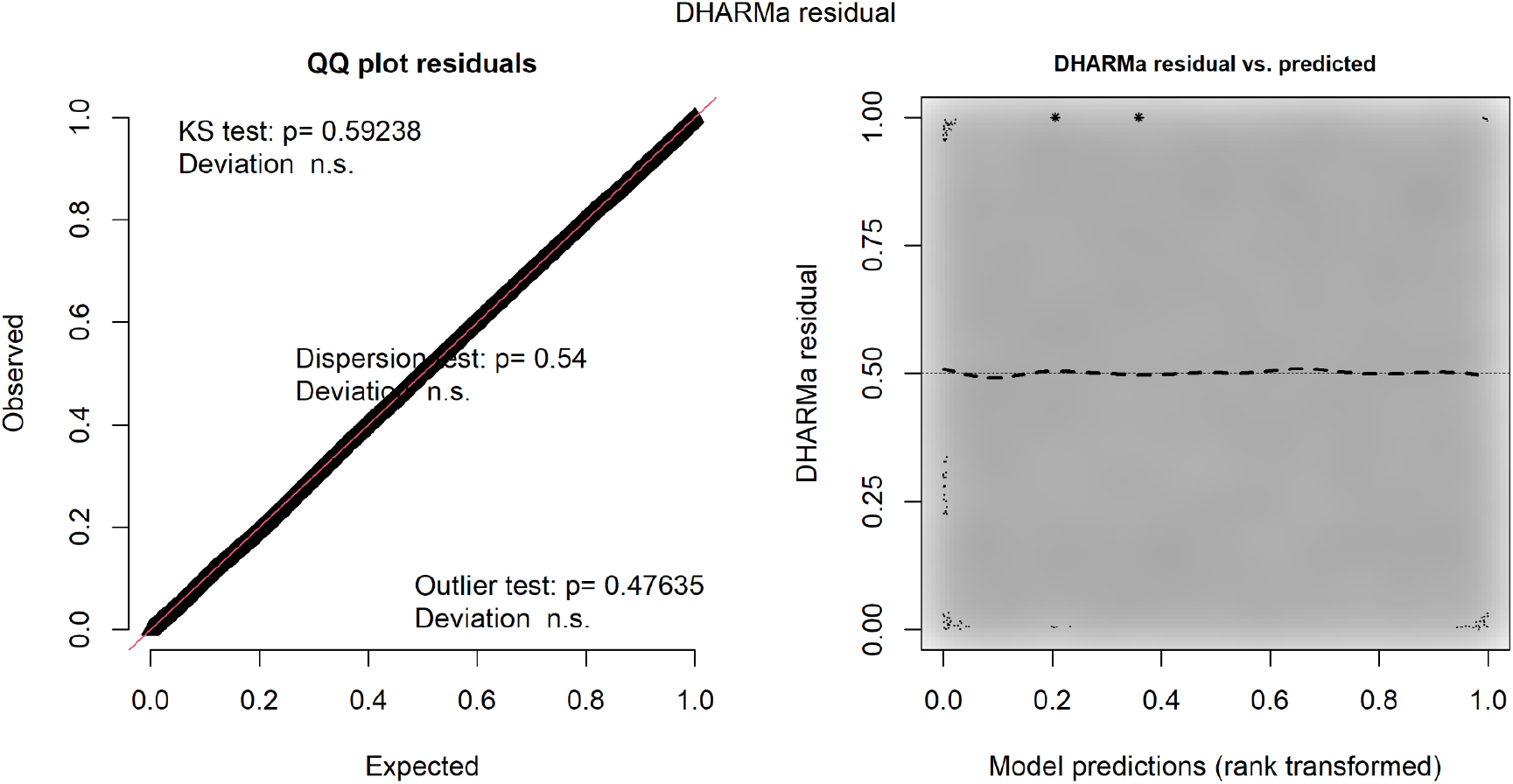
Residual diagnostics for the core binomial GLMM evaluating colonization probability. Diagnostics were generated using the “DHARMa” R package to validate the assumptions of the model. (Left) The Quantile-Quantile plot of simulated scaled residuals demonstrates perfect alignment with the expected uniform distribution. Formal goodness-of-fit tests confirm no significant deviation from the expected distribution, no overdispersion, and no significant outliers. (Right) The plot of standardized residuals against rank-transformed model predictions displays a flat, uniform dispersion along the expected horizontal quantiles. This confirms the absence of heteroscedasticity and indicates no unmodeled nonlinear patterns.

### Supplemental Tables

**Table S1.** GLMM summary for the relationship between colonization probability and model predictors (timeseries = 1,059, n = 24,707 presence and absence records, hexagons =134, species =75). Significant predictors are bolded.

| Variable | Coefficient | Std. error | Z value | P value | Explained variance component (%) |
| --- | --- | --- | --- | --- | --- |
| **Year of first record** | **-0.47** | **0.075** | **-6.31** | **< 0.0001** | **1.74** |
| CPI | 0.011 | 0.055 | 0.197 | 0.8441 | -0.33 |
| **Sources** | **0.342** | **0.026** | **13.051** | **< 0.0001** | **6.95** |
| Cumulative intensity | -0.015 | 0.042 | -0.366 | 0.7145 | 1.19 |
| **Year** | **1.602** | **0.052** | **31.057** | **< 0.0001** | **66.45** |
| **SST - annual** | **0.205** | **0.086** | **2.394** | **0.0167** | **26.2** |
| SST - climate | 0.036 | 0.076 | 0.468 | 0.6397 | -1.99 |
| **Marine area** | **-0.107** | **0.047** | **-2.268** | **0.0233** | **0.17** |
| SSS - annual | 0.04 | 0.056 | 0.719 | 0.4724 | -0.38 |
| **R-squared (marginal/conditional)** | **R<sup>2</sup>m = 0.478** | **R<sup>2</sup>c = 0.528** |  |  |  |

**Table S2.** Model selection table showing all converged model combinations. Yellow marker highlight models that can be equally considered as the most parsimonious models based on their delta AICc score (i.e., ≤ 2), balancing over- and underfit. Here we show the 10 models with the lowest AICc score out of 512 models.

| Model | Cumulative intensity | SSS | CPI | Year of first record | Marine area | SST - spatial | SST - spatiotemporal | Sources | Year | df | logLik | AICc | delta | weight |
| --- | --- | --- | --- | --- | --- | --- | --- | --- | --- | --- | --- | --- | --- | --- |
| 1 | - | - | - | -0.465 | -0.108 | - | 0.254 | 0.342 | 1.596 | 8 | -3262.8 | 6541.612 | 0 | 0.199 |
| 2 | - | 0.037 | - | -0.468 | -0.11 | - | 0.234 | 0.342 | 1.596 | 9 | -3262.57 | 6543.148 | 1.536 | 0.092 |
| 3 | -0.021 | - | - | -0.465 | -0.109 | - | 0.255 | 0.342 | 1.594 | 9 | -3262.68 | 6543.358 | 1.746 | 0.083 |
| 4 | - | - | - | -0.465 | -0.107 | 0.03 | 0.232 | 0.342 | 1.601 | 9 | -3262.72 | 6543.447 | 1.836 | 0.079 |
| 5 | - | - | 0.018 | -0.467 | -0.108 | - | 0.251 | 0.341 | 1.598 | 9 | -3262.75 | 6543.509 | 1.898 | 0.077 |
| 6 | - | - | - | -0.463 | - | - | 0.258 | 0.338 | 1.598 | 7 | -3265.35 | 6544.707 | 3.095 | 0.042 |
| 7 | - | 0.044 | - | -0.469 | -0.107 | 0.042 | 0.2 | 0.342 | 1.603 | 10 | -3262.41 | 6544.834 | 3.222 | 0.04 |
| 8 | -0.019 | 0.035 | - | -0.468 | -0.11 | - | 0.236 | 0.342 | 1.595 | 10 | -3262.46 | 6544.934 | 3.322 | 0.038 |
| 9 | - | 0.036 | 0.014 | -0.47 | -0.109 | - | 0.232 | 0.342 | 1.598 | 10 | -3262.54 | 6545.083 | 3.471 | 0.035 |
| 10 | -0.018 | - | - | -0.465 | -0.107 | 0.025 | 0.237 | 0.342 | 1.599 | 10 | -3262.62 | 6545.249 | 3.638 | 0.032 |

**Table S3.** Relative importance of predictors based on the sum of AIC weights (SW). The SW for a predictor is the sum of the AIC Weights of all models in the candidate set that contain that variable. It ranges from 0 to 1 and represents the evidence for a variable’s inclusion in the most probable set of models. In other words, a predictor with a SW of 1 is highly important as it appears in all the best-supported models (e.g., ‘Year’ appears in models that collectively account for 100% of the AIC weight).

| **Predictor** | **SW** |
| --- | --- |
| Sources | 1 |
| Year | 1 |
| Year of first record | 1 |
| SST - annual | 0.977 |
| Marine area | 0.819 |
| SSS - annual | 0.327 |
| SST - climate | 0.309 |
| Cumulative intensity | 0.29 |
| CPI | 0.279 |

**Table S4.** Pre-establishment phase. GLMM summary for the relationship between colonization probability and model predictors for the first three successive movements (i.e., t_0_,t_1_,t_2_ by Vagenas et al. 2024), where each species was observed (timeseries = 117, n = 2,634, presence and absence records, hexagons = 22, species = 45).

| Pre-establishment phase |  |  |  |  |  |
| --- | --- | --- | --- | --- | --- |
| Variable | Coefficient | Std. error | Z value | P value | Explained variance component (%) |
| **Year of first record** | **-1.632** | **0.21** | **-7.774** | **< 0.0001** | **14** |
| CPI | -0.139 | 0.195 | -0.715 | 0.4746 | 0.07 |
| **Sources** | **0.573** | **0.096** | **5.967** | **< 0.0001** | **13.83** |
| Cumulative intensity | 0.13 | 0.128 | 1.018 | 0.3088 | 0.19 |
| **Year** | **2.67** | **0.302** | **8.834** | **< 0.0001** | **64.77** |
| SST - annual | -0.188 | 0.275 | -0.682 | 0.4952 | 6.35 |
| SST - climate | 0.416 | 0.23 | 1.808 | 0.0706 | 0.42 |
| Marine area | 0.091 | 0.168 | 0.543 | 0.5872 | 0.04 |
| SSS | 0.039 | 0.186 | 0.209 | 0.8346 | 0.32 |
| **R-squared (marginal/conditional)** | **$R^2m = 0.69$** | **$R^2c = 0.729$** |  |  |  |

**Table S5.**
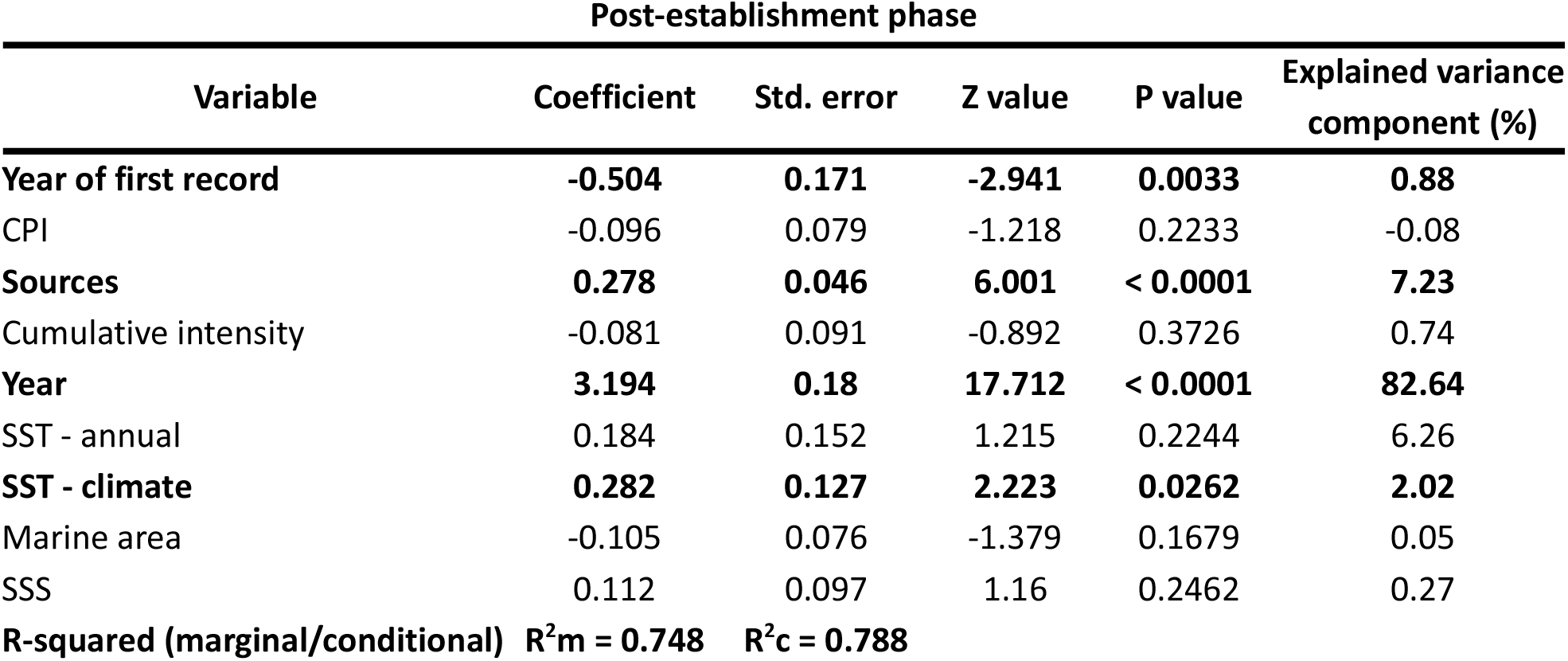
Post-establishment phase. GLMM summary for the relationship between colonization probability and model predictors for the fourth to the maximum successive record (i.e., t_3_,t_4_…t_max_) where each species was observed (timeseries = 315, n = 8,532 presence and absence records, hexagons = 95, species = 31).

**Table S6.**
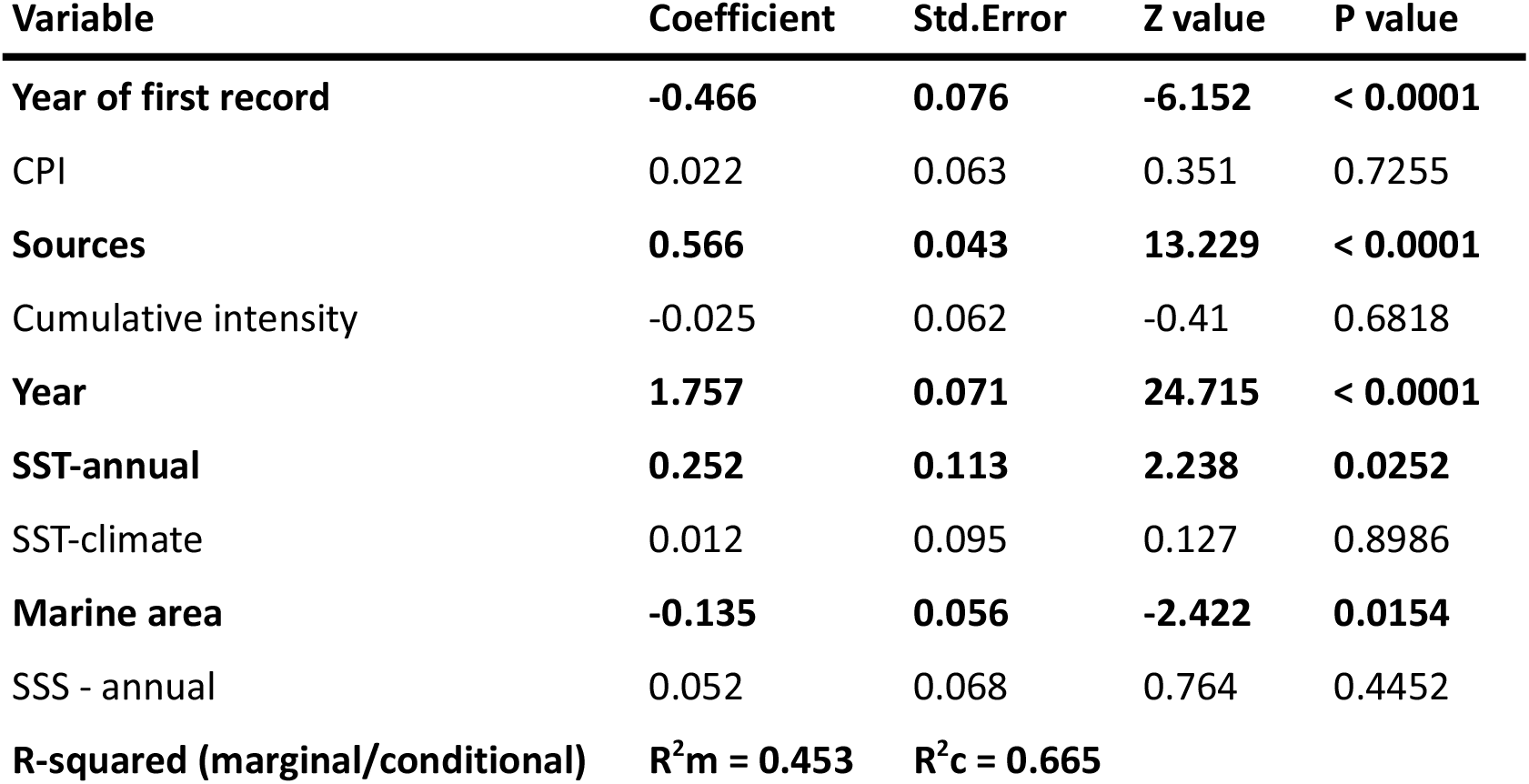
Sensitivity analysis for spatio-temporal pseudoreplication. Summary of the binomial GLMM evaluating colonization probability while explicitly accounting for unmeasured, transient local anomalies. In addition to the core model predictors and baseline random intercepts for spatial location (hexagon) and taxonomy (species), this model incorporates a crossed random effect for the interaction between space and time (hexagon:year). This interaction structure ensures that concurrent observations of multiple species within the exact same hexagon and year do not artificially inflate the significance of the fixed-effect estimates (n resence and absence records = 24,707; hexagons = 134; species = 75).

**Table S7.** Canal enlargement sensitivity GLMM summary. (timeseries = 1,059, n = 24,707 presence and absence records, hexagons = 134, species = 75).

| **Variable** | **Coefficient** | **Std.Error** | **Z value** | **P value** |
| --- | --- | --- | --- | --- |
| **Year of first record** | **-0.48** | **0.076** | **-6.307** | **< 0.0001** |
| CPI | 0.012 | 0.056 | 0.216 | 0.8289 |
| **Sources** | **0.344** | **0.026** | **12.971** | **< 0.0001** |
| Cumulative intensity | 0.014 | 0.042 | 0.327 | 0.7434 |
| **Year** | **1.897** | **0.097** | **19.519** | **< 0.0001** |
| **SST-annual** | **0.195** | **0.087** | **2.247** | **0.0246** |
| SST-climate | 0.054 | 0.078 | 0.691 | 0.4896 |
| **Marine area** | **-0.108** | **0.048** | **-2.227** | **0.0259** |
| **Canal area** | **-0.333** | **0.093** | **-3.581** | **0.0003** |
| SSS - annual | 0.038 | 0.057 | 0.659 | 0.5099 |
| **R-squared (marginal / conditional)** | **R<sup>2</sup>m = 0.477** | **R<sup>2</sup>c = 0.531** |  |  |

**Table S8.** The bimodal oscillating system (BiOS) sensitivity GLMM summary. (timeseries = 1,038, n = 19,443 presence and absence records, hexagons = 134, species = 75).

| **Variable** | **Coefficient** | **Std.Error** | **Z value** | **P value** |
| --- | --- | --- | --- | --- |
| **Year of first record** | **-0.461** | **0.075** | **-6.11** | **< 0.0001** |
| CPI | 0.015 | 0.057 | 0.26 | 0.7952 |
| **Sources** | **0.36** | **0.029** | **12.501** | **< 0.0001** |
| Cumulative intensity | 0.04 | 0.044 | 0.917 | 0.3593 |
| **Year** | **1.435** | **0.048** | **30.151** | **< 0.0001** |
| **SST-annual** | **0.243** | **0.088** | **2.762** | **0.0057** |
| SST-climate | 0.022 | 0.079 | 0.284 | 0.7762 |
| **Marine area** | **-0.113** | **0.049** | **-2.316** | **0.0206** |
| SSS-annual | 0 | 0.058 | -0.001 | 0.9989 |
| **BiOS phase - cyclonic** | **-0.309** | **0.074** | **-4.17** | **< 0.0001** |
| **R-squared (marginal / conditional)** | **R<sup>2</sup>m = 0.422** | **R<sup>2</sup>c = 0.48** |  |  |

**Table S9.** Thermal range sensitivity GLMM summary. (timeseries = 951, n = 22,206 presence and absence records, hexagons = 134, species = 66).

| **Variable** | **Coefficient** | **Std.Error** | **Z value** | **P value** |
| --- | --- | --- | --- | --- |
| **Year of first record** | **-0.456** | **0.08** | **-5.683** | **< 0.0001** |
| CPI | -0.002 | 0.059 | -0.036 | 0.9711 |
| **Sources** | **0.355** | **0.028** | **12.779** | **< 0.0001** |
| Cumulative intensity | -0.022 | 0.045 | -0.493 | 0.6224 |
| **Year** | **1.627** | **0.055** | **29.511** | **< 0.0001** |
| **SST-annual** | **0.177** | **0.092** | **1.928** | **0.0538** |
| SST-climate | 0.064 | 0.082 | 0.774 | 0.4388 |
| **Marine area** | **-0.12** | **0.05** | **-2.402** | **0.0163** |
| SSS - annual | 0.021 | 0.059 | 0.349 | 0.7268 |
| Thermal range | 0.046 | 0.094 | 0.494 | 0.6214 |
| **R-squared (marginal / conditional)** | **R<sup>2</sup>m = 0.481** | **R<sup>2</sup>c = 0.534** |  |  |

**Table S10.**
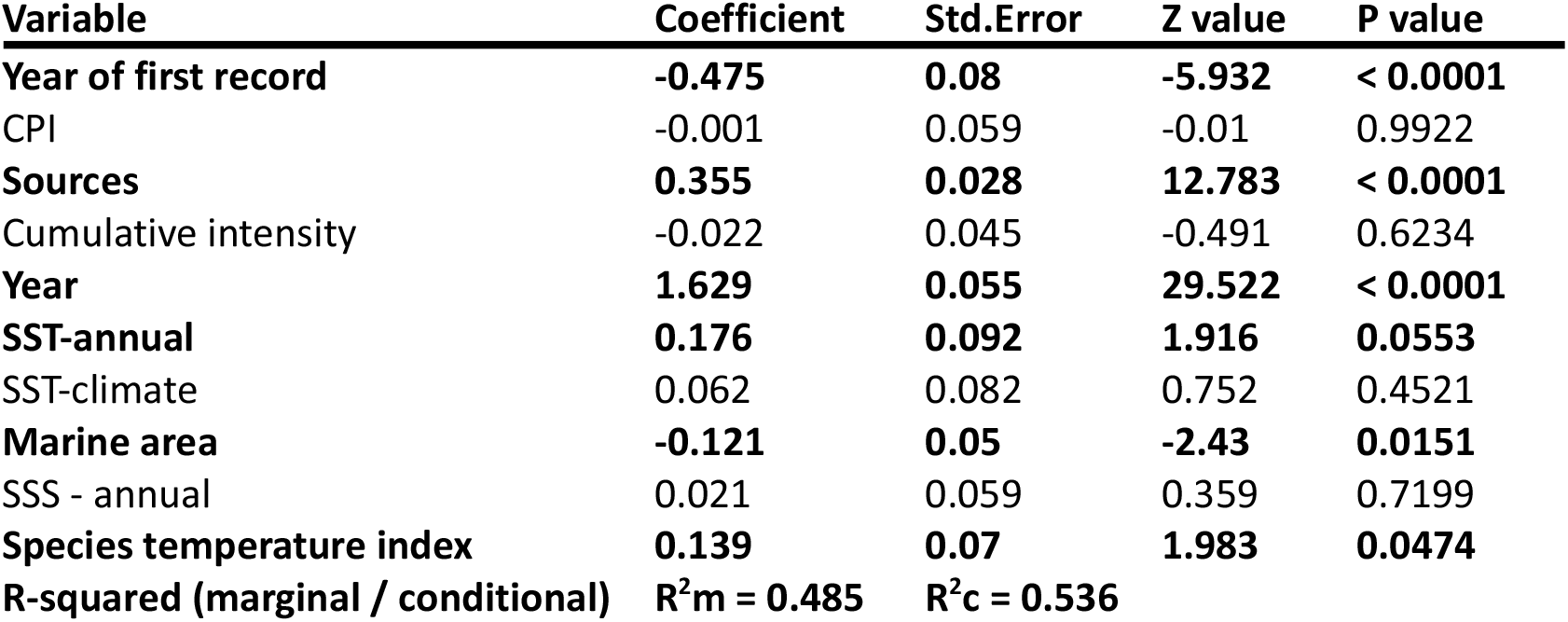
Species temperature index sensitivity GLMM summary. (timeseries = 951, n = 22,206 presence and absence records, hexagons = 134, species = 66).

| **Variable** | **Coefficient** | **Std.Error** | **Z value** | **P value** |
| --- | --- | --- | --- | --- |
| **Year of first record** | **-0.475** | **0.08** | **-5.932** | **< 0.0001** |
| CPI | -0.001 | 0.059 | -0.01 | 0.9922 |
| **Sources** | **0.355** | **0.028** | **12.783** | **< 0.0001** |
| Cumulative intensity | -0.022 | 0.045 | -0.491 | 0.6234 |
| **Year** | **1.629** | **0.055** | **29.522** | **< 0.0001** |
| **SST-annual** | **0.176** | **0.092** | **1.916** | **0.0553** |
| SST-climate | 0.062 | 0.082 | 0.752 | 0.4521 |
| **Marine area** | **-0.121** | **0.05** | **-2.43** | **0.0151** |
| SSS - annual | 0.021 | 0.059 | 0.359 | 0.7199 |
| **Species temperature index** | **0.139** | **0.07** | **1.983** | **0.0474** |
| **R-squared (marginal / conditional)** | **R<sup>2</sup>m = 0.485** | **R<sup>2</sup>c = 0.536** |  |  |

**Table S11.**
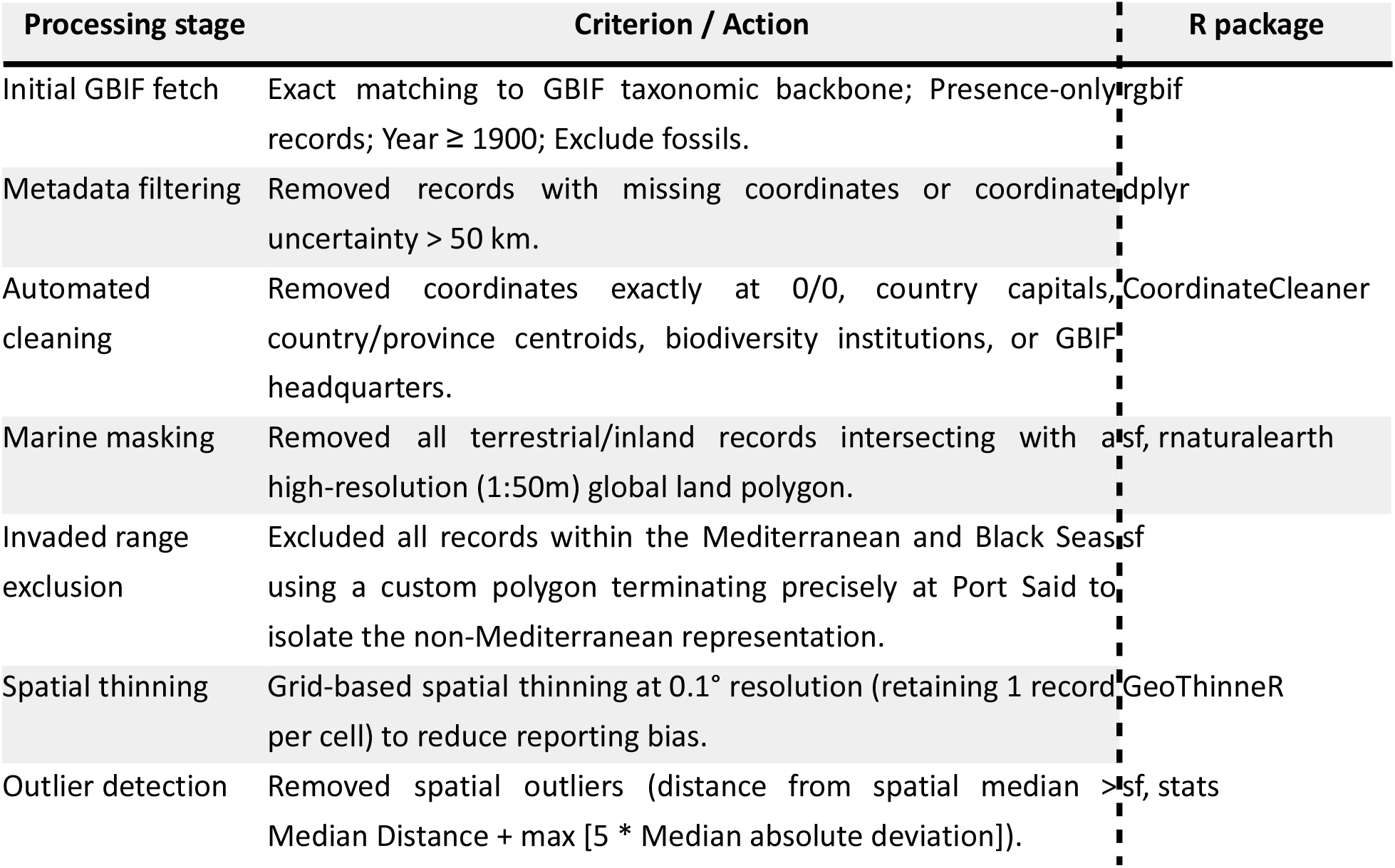
Sequential filtering and processing criteria applied to GBIF occurrence records to estimate the species temperature index.

**Table S12.**
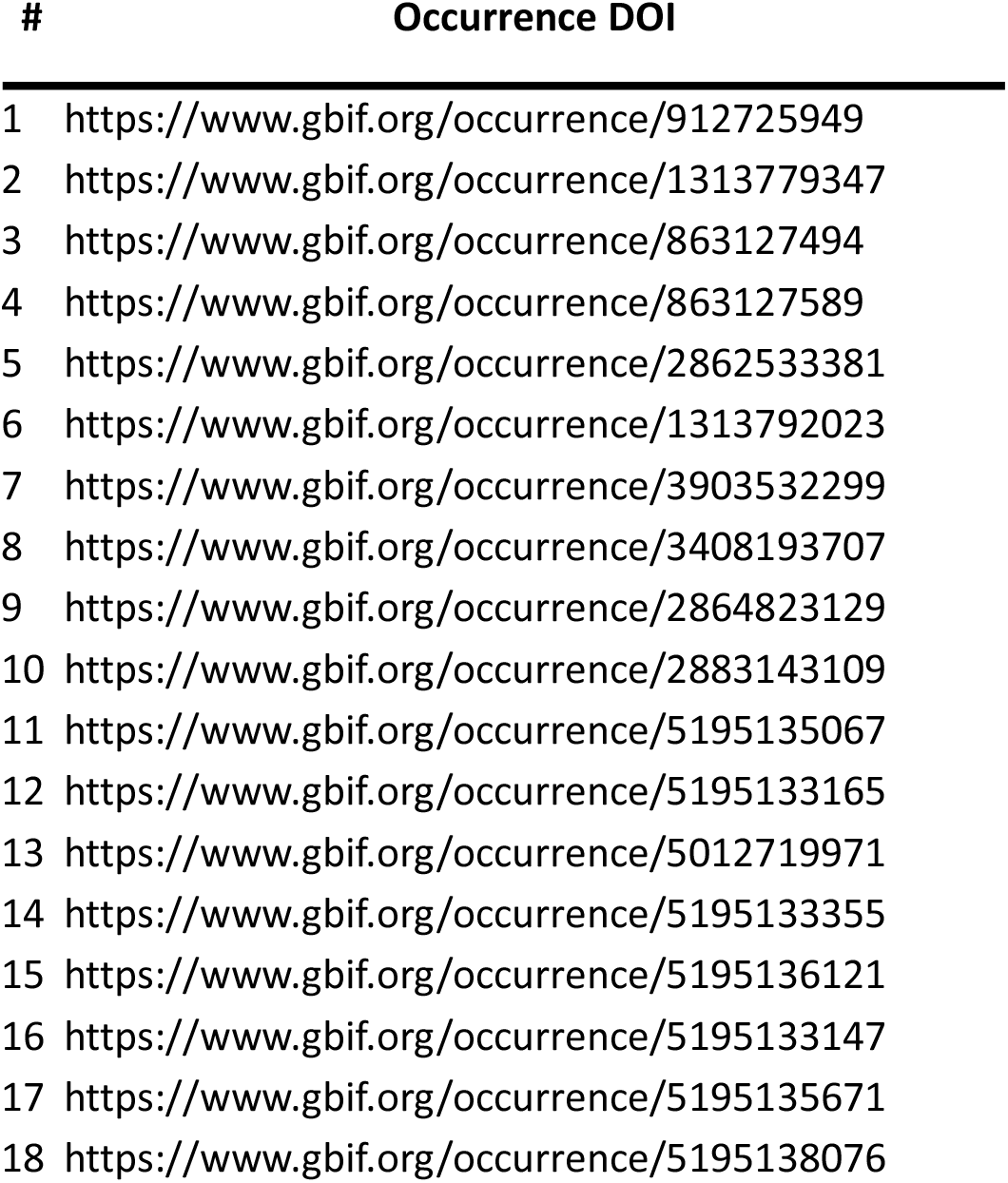
Supplemented GBIF records.

| # | Occurrence DOI |
| --- | --- |
| 1 | https://www.gbif.org/occurrence/912725949 |
| 2 | https://www.gbif.org/occurrence/1313779347 |
| 3 | https://www.gbif.org/occurrence/863127494 |
| 4 | https://www.gbif.org/occurrence/863127589 |
| 5 | https://www.gbif.org/occurrence/2862533381 |
| 6 | https://www.gbif.org/occurrence/1313792023 |
| 7 | https://www.gbif.org/occurrence/3903532299 |
| 8 | https://www.gbif.org/occurrence/3408193707 |
| 9 | https://www.gbif.org/occurrence/2864823129 |
| 10 | https://www.gbif.org/occurrence/2883143109 |
| 11 | https://www.gbif.org/occurrence/5195135067 |
| 12 | https://www.gbif.org/occurrence/5195133165 |
| 13 | https://www.gbif.org/occurrence/5012719971 |
| 14 | https://www.gbif.org/occurrence/5195133355 |
| 15 | https://www.gbif.org/occurrence/5195136121 |
| 16 | https://www.gbif.org/occurrence/5195133147 |
| 17 | https://www.gbif.org/occurrence/5195135671 |
| 18 | https://www.gbif.org/occurrence/5195138076 |

**Table S13.** Variance Inflation Factor (VIF) summary table for the core model presented in Table S1. All VIF values remain below the threshold of 5, supporting the absence of multicollinearity

| **Term** | **VIF** | **Tolerance** |
| --- | --- | --- |
| Year of first record | 1.031 | 0.970 |
| CPI | 1.095 | 0.913 |
| Sources | 1.091 | 0.917 |
| Cumulative intensity | 1.043 | 0.959 |
| Year | 1.137 | 0.879 |
| SST - annual | 2.978 | 0.336 |
| SST - climate | 2.374 | 0.421 |
| Marine area | 1.025 | 0.976 |
| SSS - annual | 1.471 | 0.680 |

### Supplemental Methods

#### Species selection

Out of 205 non-indigenous fish species currently present in the Mediterranean Sea (171 by 2020 as reported by (*1*) and updated with 34 new species arrivals until 2023 as documented in the recent literature), 130 are assumed to be Lessepsians, i.e., species of Indo-Pacific origin arriving progressively in the Mediterranean via the Suez Canal with their own capabilities (*2*). Of these, 80 non-indigenous fish species have arrived in the period 1987-2023 and only 51 have established viable populations, albeit 5 of them with very localized distribution. Our dataset thus includes records of the 46 widespread established species introduced in the period 1987-2023, with the addition of 29 established non-indigenous species which were introduced before 1987 but spread from the eastern Mediterranean (as delineated for the purposes of the EU Marine Strategy Framework Directive [MSFD] – (*3*) in new MSFD areas in the period 1987-2023 eg. *Hyporamphus affinis*.

Established non-indigenous species excluded from our analysis belong to three categories: A) species introduced before 1987 and established in the eastern Mediterranean, that have not spread in neighbouring MSFD areas e.g. *Upeneus moluccensis*, *Cynoglossus sinusarabici*, *Lutjanus argentimaculatus*; B) species introduced in the 1987-2023 period but with local distribution, e.g. *Ambassis dussumieri* (2021, Israel, Gaza port - *4*), *Hazeus ingressus* (2014-15, S. Turkey - *5*); *Stolephorus indicus* (2015, Israel - *6*); *Cyclichthys spilostylus* (1993, Israel - *7*); *Tetrosomus gibbosus* (1987, Israel - *8*) those with unresolved taxonomic status such as *Abudefduf cf. saxatilis/vaigiensis/troschelii*.

It should be noted that for a small number of species, their first Mediterranean records were attributed to pathways other than the corridor (i.e., the Suez Canal); these were excluded from our analysis, which only includes subsequent records indicating additional introduction events through the Suez Canal. These species are: (1) *Fistularia petimba* Lacepède, 1803, native to the tropical Atlantic and Indo-Pacific, was first observed in Spain in 1988 (*9*) and assumed to be a range expansion; later records in the Eastern Mediterranean after 2016 (*10*) and references therein) confirmed the species as a new Lessepsian migrant in the region. (2) *Pomadasys stridens* (Forsskål, 1775) of Indo-Pacific origin was first reported in 1968 from the Ligurian Sea (*11*), presumably as a result of secondary spread as a stowaway from a yet undetected Lessepsian introduction (*12*). This was confirmed by later findings in Israel and Egypt in 1971 and 1973 respectively (*13*), and subsequently many other eastern Mediterranean locations. (3) *Diodon hystrix* Linnaeus, 1758, with a circumtropical native distribution, has been observed several times since 1953 in the central and western Mediterranean, where it is considered a vagrant or range-expanding species (*14*) and references therein); nevertheless, more recent records from the Levant are suggestive of a separate introduction event via the Suez Canal or even, possibly, the aquarium trade in the region (*15*).

